# G-quadruplexes rescuing protein folding

**DOI:** 10.1101/2022.09.09.507295

**Authors:** Ahyun Son, Veronica Huizar Cabral, Theodore J Litberg, Scott Horowitz

## Abstract

Maintaining the health of the proteome is a critical cellular task. Recently, we found G-quadruplex (G4) nucleic acids are especially potent at preventing protein aggregation *in vitro* and could at least indirectly improve the protein folding environment of *E. coli*. However, the roles of G4s in protein folding were not yet explored. Here, through *in vitro* protein folding experiments, we discover that G4s can accelerate protein folding by rescuing kinetically trapped intermediates to both native and near-native folded states. Time-course folding experiments in *E. coli* further demonstrate that these G4s primarily improve protein folding quality in *E. coli* as opposed to preventing protein aggregation. The ability for a short nucleic acid to rescue protein folding opens up the possibility of nucleic acids and ATP-independent chaperones to play considerable roles in dictating the ultimate folding fate of proteins.

## Introduction

Proteostasis is maintained through the action of a variety of proteins, particularly molecular chaperones and degradation machineries. Chaperones interact with unfolded, partially folded, or misfolded proteins to prevent aggregation, assist in protein folding, and/or target proteins for degradation (Balch et al., 2008; Hartl et al., 2011; Klaips et al., 2018; Powers et al., 2009). ATP-independent chaperones (holdases) act as a first line of defense against toxic aggregation by binding proteins as they misfold or unfold (Eisenhardt, 2013; Haslbeck and Vierling, 2015). At the cessation of stress, holdases transfer their clients to ATP-dependent chaperones (foldases) that can both prevent aggregation and directly help proteins to fold (Eisenhardt, 2013; Gray et al., 2014; Hartl *et al*., 2011). In addition to traditional protein chaperones, several other molecules have been shown to play major roles in protein folding and aggregation. One such group of non-proteinaceous chaperones are nucleic acids, which were first implicated in protein aggregation over 20 years ago when they were found in the protein aggregates in the brains from AD patients (Ginsberg et al., 1997; Ginsberg et al., 1998).

More recently, the role of nucleic acids in proteostasis has begun to receive attention (Aarum et al., 2020; Alriquet et al., 2019; Audas et al., 2012; Bounedjah et al., 2014; Cheung et al., 2011; Cheung et al., 2013; Cordeiro et al., 2001; Dinkel et al., 2015; Frottin et al., 2019; Kampers et al., 1996; Liu and Zhang, 2011; Maharana et al., 2018; Mateju et al., 2017; Rentzeperis et al., 1999; Sanchez de Groot et al., 2019; Silva and Cordeiro, 2016; Yin et al., 2009; Yin et al., 2010). For example, ∼10% of the proteome requires RNA for maintaining solubility, including many proteins that are not considered to be RNA binding proteins (Aarum *et al*., 2020). Moreover, stress granules, which are liquid-liquid phase separated condensates where proteins retreat under stress for protection, contain high levels of RNA (Maharana *et al*., 2018; Mateju *et al*., 2017; Namkoong et al., 2018). Nucleic acids maintain fluidity and solubility of stress granules (Maharana *et al*., 2018; Mateju *et al*., 2017), and in some cases are responsible for binding and guiding proteins to these condensates under stress (Alriquet *et al*., 2019; Bounedjah *et al*., 2014). These and other studies strongly suggest that RNAs constitute important players of multiple mechanisms in proteostasis. However, the precise roles of nucleic acids in proteostasis are still unclear.

We recently demonstrated that certain nucleic acids serve as powerful modulators of protein aggregation, oligomerization, and protein folding (Begeman et al., 2020; Docter et al., 2016; Horowitz and Bardwell, 2016; Litberg et al., 2020). Intriguingly, we discovered that G4-containing sequences are especially potent as protein chaperones. We found that G4s prevent aggregation *in vitro* more efficiently on a weight basis than known chaperone proteins, and an order of magnitude more efficiently than bulk DNA (Begeman *et al*., 2020). Moreover, we demonstrated that many G4s could also promote a healthy protein folding environment in cells, although whether this was due to affecting protein folding directly was unclear at the time. Of note, unfolded nucleic acids did not show these effects *in vitro* or in cells, suggesting these properties were special to G4s (Begeman *et al*., 2020).

G4s are nucleic acids that contain polyG stretches that form base tetrads. G4s can combine with other nucleic acid structural elements to create highly complex nucleic acid structures (Banco and Ferre-D’Amare, 2021; Reddy Sannapureddi et al., 2020) and have previously been characterized to have roles in regulating transcription, translation, and splicing (Varshney et al., 2020). Although the full extent of G4 formation is not yet known in the cell, they have been identified in DNA, mRNA, and in the large rRNA extensions of ribosomes as well as in tRNA that is cleaved under stress conditions to create G4-forming tiRNA (Ivanov et al., 2014; Jansson et al., 2019; Mestre-Fos et al., 2020; Shao et al., 2020).

Foldases are typically considered to aid proteins in folding in three major ways. In the case of Hsp60s (such as *E. coli* GroEL), there is significant evidence that these chaperones can directly catalyze protein folding, which is hypothesized to occur through a combination of enclosure and electrostatic effects (Chakraborty et al., 2010; Georgescauld et al., 2014; Singh et al., 2020), as well as to increase apparent folding speed by isolating the protein from off-pathway aggregation (Tyagi et al., 2011). In Hsp60s as well as other foldases, there is also evidence of unfolding mechanisms in which the chaperone helps to unfold kinetically trapped misfolded states to allow another chance at folding (Priya et al., 2013a; Priya et al., 2013b; Saibil, 2013; Thirumalai et al., 2020). Work with GroEL has shown multiple mechanisms to be in play for its ability to modulate of protein folding (Chakraborty *et al*., 2010; Georgescauld *et al*., 2014; Koldewey et al., 2017; Priya *et al*., 2013b; Singh *et al*., 2020; Thirumalai *et al*., 2020), but it remains a controversial topic (Ambrose et al., 2015; Tyagi *et al*., 2011). With ATP-independent holdases, effects on folding have previously been found to be primarily passive (Horowitz et al., 2018), but it remains possible that there are direct effects on protein folding.

Before finding the potency of G4s as chaperones, we investigated the ability of disordered and unfolded ssRNA to affect protein folding. We examined the ability of polyU RNA to help the folding of luciferase. In these experiments, polyU had no effect on the refolding of luciferase on its own, but could serve as a holdase during stress that transferred the partially folded protein to Hsp70 for refolding. As a result, in the presence of polyU RNA and Hsp70, a much greater amount of luciferase was refolded than with just Hsp70 (Docter *et al*., 2016). We thus hypothesized that nucleic acids function as holdases, similar to known ATP-independent chaperones. Unexpectedly, our work here reveals that G4s accelerate protein folding by rescuing kinetically trapped intermediates, a property previously thought to be predominantly performed by ATP-dependent foldases. Continuing experiments in *E. coli* shows that the effects of chaperone nucleic acids are not protease, solubility, or expression level-dependent, but promote protein folding.

## Results

### G4-containing DNA rescues protein folding of TagRFP675 *in vitro*

To investigate whether G4s could not only prevent aggregation but also affect protein folding, we checked whether G4s we previously found that had good holdase activity (such as Seq576) (Begeman *et al*., 2020) could potentially affect protein refolding *in vitro*. To test this question, we purified TagRFP675, and performed *in vitro* refolding assays. In these assays, TagRFP675 is chemically denatured, and then rapidly diluted into a native buffer to allow refolding in the presence or absence of G4-containing chaperones. We monitored the refolding of TagRFP675 by the appearance of its fluorescence, which depends almost solely on its chromophore formation and being in the native state (Craggs, 2009). At room temperature, we found that a small amount of protein renatured quickly, followed by a continued slow increase in folded protein (**Figure 1A**). Adding Seq576 (Begeman *et al*., 2020) caused a marked increase in the speed of protein refolding (**Figure 1B**). A second G4 of known structure, LTR-III (Butovskaya et al., 2018), also instigated substantial protein folding **(Figure 1C)**. Contrasting the effects of the G4s, using a ssDNA sequence of the same length but containing no G4 structure (Seq42) (**Figure 1D**) caused no increase in the speed of refolding. This experiment suggests that the G4 aids in the folding of the protein, unlike an unfolded nucleic acid that has no effect on folding.

**Figure 1.**
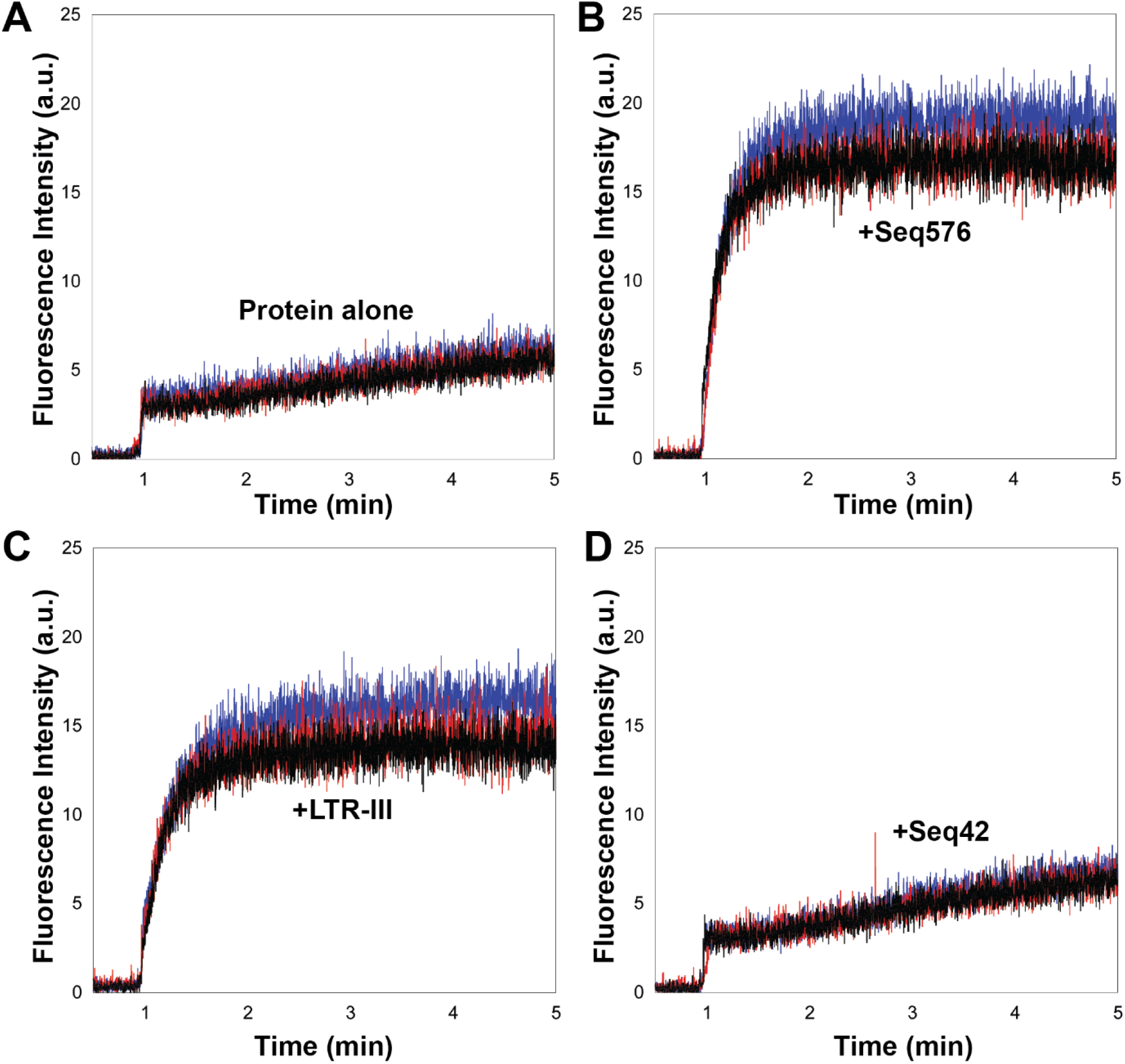
Fluorescence recovery of TagRFP675 in the presence of nucleic acids containing G4 structures *in vitro*: TagRFP675 was denatured in 6M guanidine-hydrochloride (Gu-HCl) for 11 hours at 23°C. Denatured protein was diluted to 0.5 μM into potassium phosphate and the increase in fluorescence intensity was measured over time in the absence of nucleic acids (A) and in the presence of 1 μM of Seq576 (B), LTR-III (C), and Seq42 (D). The absence of nucleic acids and the addition of Seq42 served as the negative controls and LTR-III and Seq576 contain G4 structures. Three triplicate measurements are shown with the separate colors.

The difficulty of TagRFP675 to refold on its own in this experiment is consistent with previous literature on the folding of fluorescent proteins. It has been well established that fluorescent β-barrel proteins display a hysteresis in folding, in which returning to the native state after chemical denaturation occurs on the timescale of days (Andrews et al., 2009; Fukuda et al., 2000; Hsu et al., 2009; Huang et al., 2007; Stepanenko et al., 2004). This very slow refolding rate is due to a rugged folding landscape, and particularly the formation of kinetically trapped intermediates that escape to the native state very slowly (Andrews *et al*., 2009; Hsu *et al*., 2009; Huang *et al*., 2007; Stepanenko *et al*., 2004). These intermediates have β-barrel formation but appear to be structurally different primarily in the lid region in the case of superfolder GFP (Andrews *et al*., 2009). We therefore hypothesized that TagRFP675 could similarly be trapped in intermediate states and unable to refold without the presence of a chaperone to assist transitioning from the kinetically trapped intermediate to the native state.

To directly test the kinetically-trapped intermediate hypothesis, we reasoned that by increasing the refolding temperature, TagRFP675 would overcome the kinetic folding barrier more quickly without a chaperone (Dill and Bromberg, 2003). In this case, we would observe an increase in the folding rate as a function of temperature. Indeed, our data showed that the rate of spontaneous refolding dramatically decreased at lower temperature and improved with increasing temperature, making the presence of the chaperone G4 almost unnecessary at 40°C (**Figure 2**). These results strongly support the hypothesis that some portion of TagRFP675 is in a kinetically trapped state that the G4 is able to rescue to the native state.

**Figure 2:**
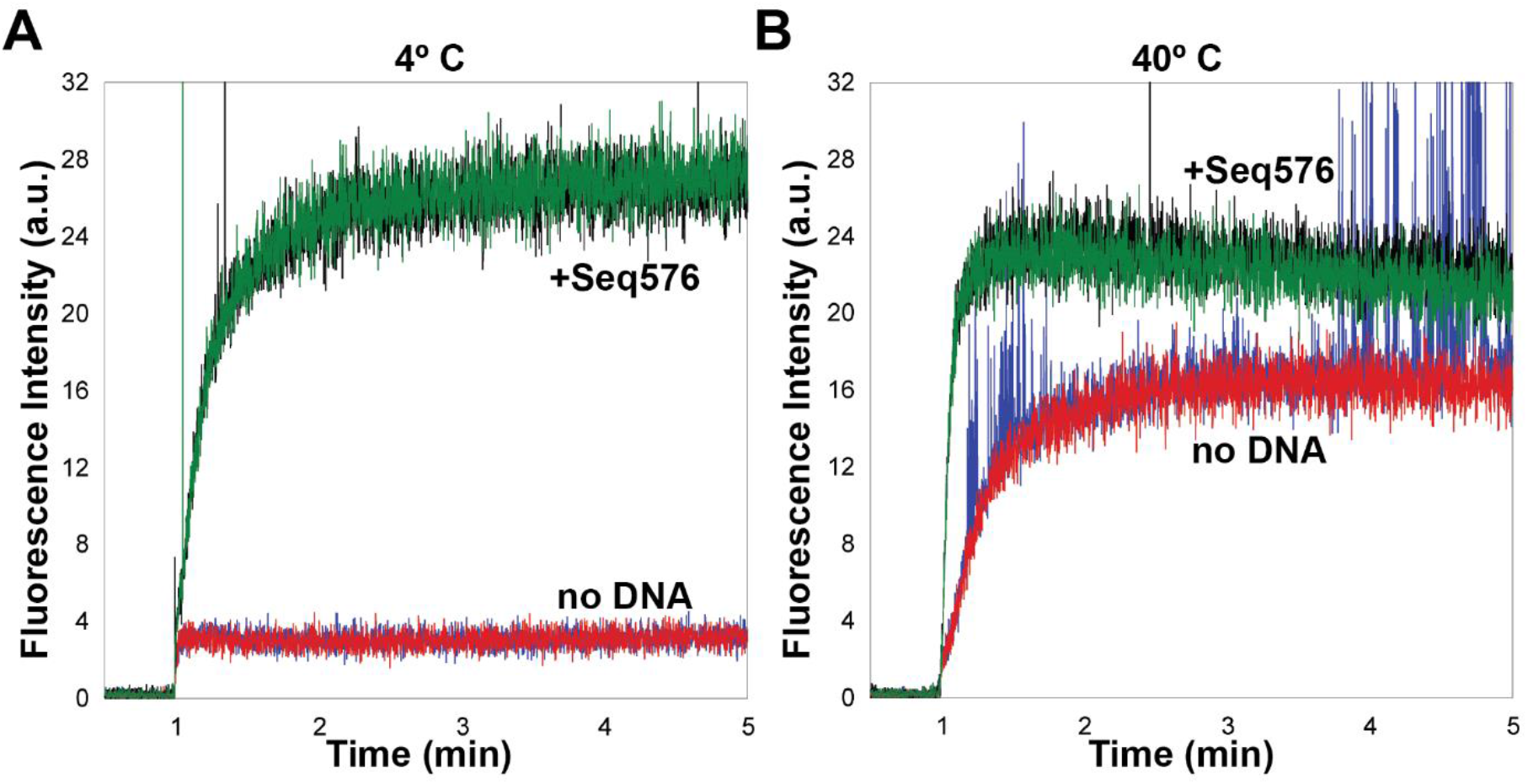
Temperature dependence of protein folding. TagRFP675 was diluted from denaturing buffer (6 M Gu-HCl) to 0.5 µM protein into refolding buffer. Addition of 1 µM Seq576 at 4°C (A) and 40°C (B). Replicates shown in separate colors. Experiments at 23°C can be seen in **Figure 1**.

We next investigated whether kinetically trapped states were being rescued by changing when the G4 was added. If kinetically trapped states were stable, we would observe the refolding regardless of when the G4 was added to the kinetically trapped protein. We therefore performed the refolding experiment as before but added Seq576 at defined times after the protein was diluted into the refolding buffer (**Figure 3**). The results showed that regardless of whether the G4 was added before or after the addition of protein, the addition of the G4 immediately prompted the refolding of TagRFP675 (**Figure 3)** to nearly the same extent. This observation suggests that there is a sub-pool of trapped intermediates that are able to overcome the kinetic barrier to the folded state by association with the G4. We also varied the concentration of Seq576 to detect the effect on protein refolding. Increasing the amount of G4 did lead to an increase in the level of protein refolding, but this effect leveled off at a small fraction of total protein folding (**Figure S1**).

**Figure 3:**
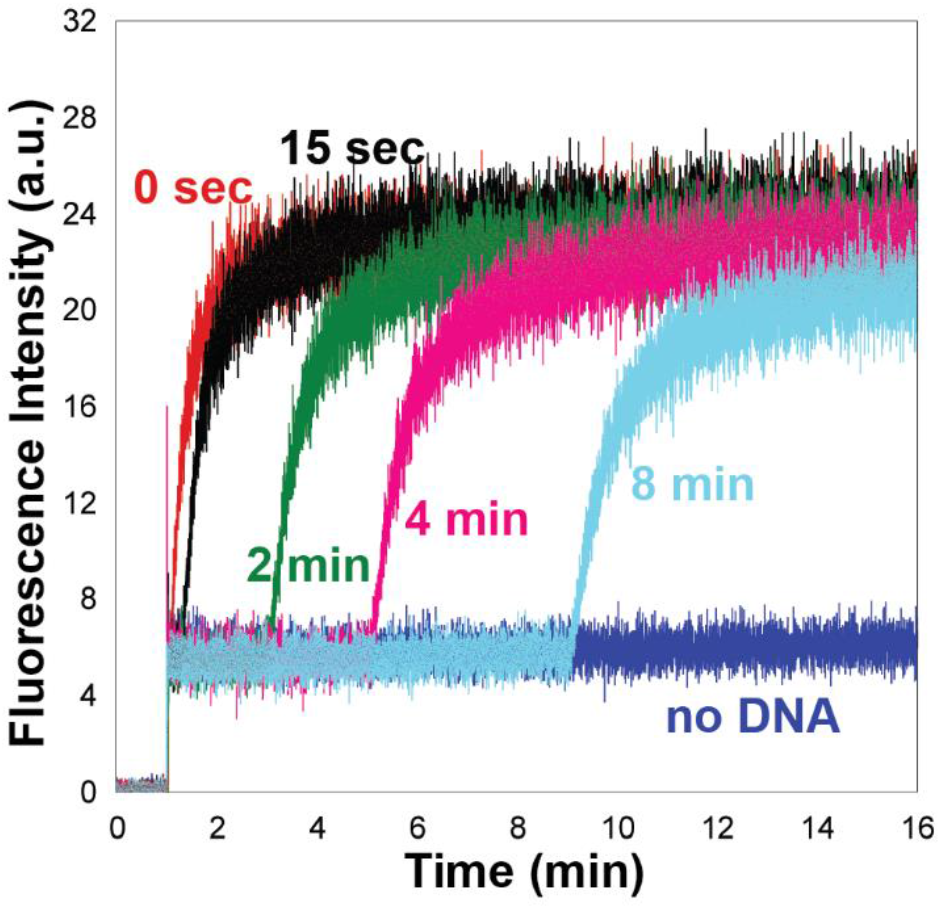
Addition of G4-containing DNA triggers refolding after dilution. Timed addition of Seq576, showing folding rescue at timepoints after TagRFP675 dilution into refolding buffer at 4°C. TagRFP675 was diluted from denaturing buffer (6 M Gu-HCl) to 0.5 µM protein into refolding buffer followed by timed addition of Seq576.

TagRFP675’s chromophore forms an intricate set of hydrogen bonds that affect its emission spectrum. Specifically, the hydroxyl group forms several hydrogen bonds that shift the emission maximum from 510 nm to 675 nm (Piatkevich et al., 2013). By also monitoring 510 nm during refolding, we could therefore separately observe the protein in an intermediate state that is distinct from the native state. Monitoring at 510 nm in the absence of Seq576, we observed roughly an order of magnitude increase in fluorescence relative to that observed at 675 nm, suggesting that a considerably larger fraction of the protein collapses into a non-native fluorescent intermediate than to the native state (**Figure 4A**). Performing this same experiment in the presence of Seq576 caused a considerable increase in the fast formation of this non-native intermediate over the case without G4 (**Figure 4A**). This observation suggests that the G4 is able to promote the formation of both fluorescent non-native intermediates as well as to rescue a smaller portion of kinetically trapped intermediates to the native state. Furthermore, in both cases the amount of protein in this partially folded state observed at 510 nm decreases slowly over time that roughly corresponds with the slow increase in fluorescence at 675 nm, suggesting that there is a slow conversion from this intermediate state to the native state (**Figure 4A**). Like with the native state, delayed addition of the G4 also rescues folding to the intemediate state (**Figure 4B**). Overall, these results show that G4s can rescue protein folding from intermediate states.

**Figure 4:**
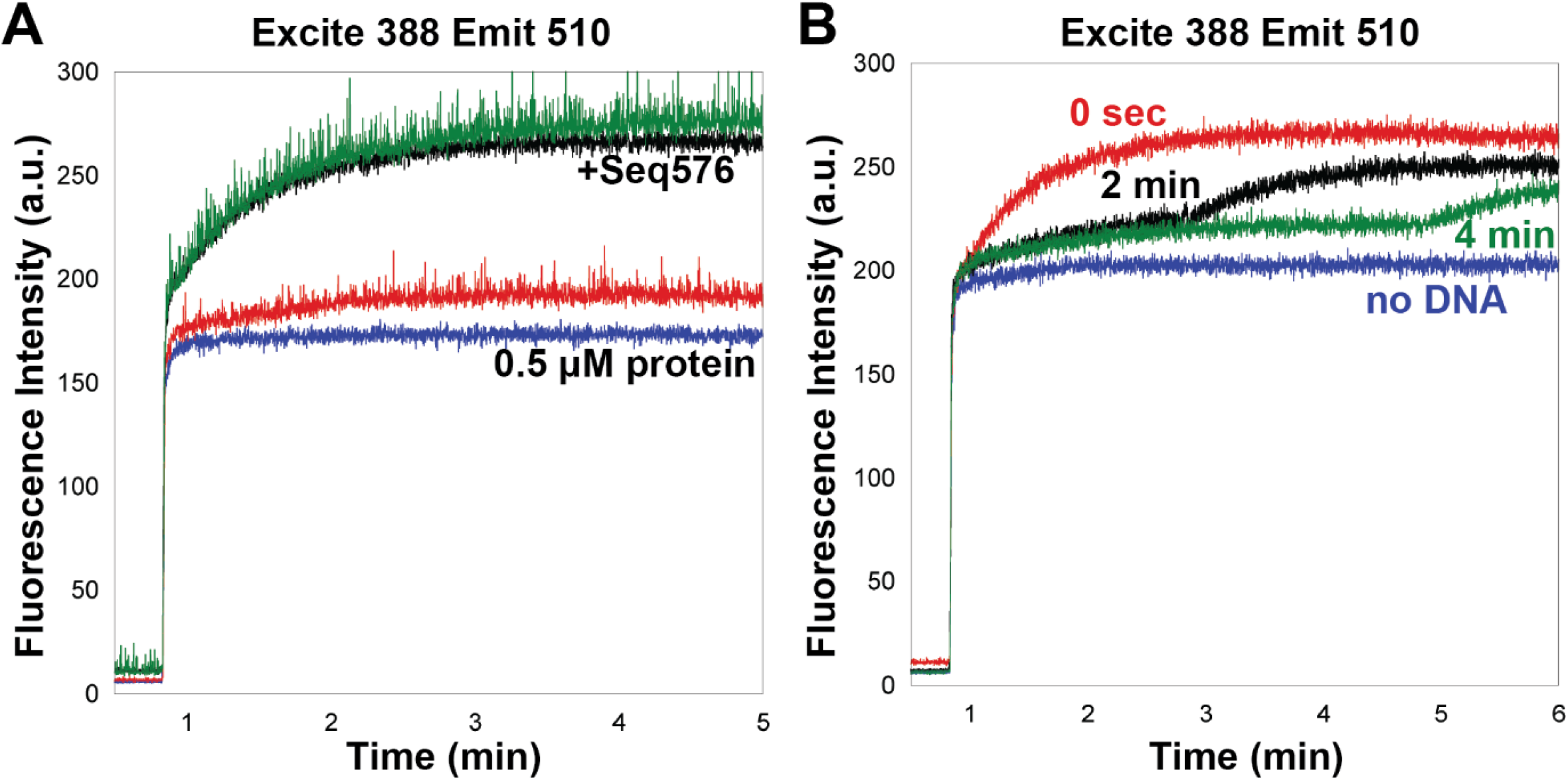
Observing folding to intermediate state. TagRFP675 was diluted from denaturing buffer (6 M Gu-HCl) to 0.5 µM protein into refolding buffer in the presence of 1 µM Seq576 at 23°C and observed at 510 nm emission at 23°C either before dilution (A) or with timed addition of Seq576 (B).

### G4 enhances proper folding of TagRFP675 in cells

Having seen an effect of promoting protein folding *in vitro*, we next examined the effects of a G4 on folding in cells. Previously, we showed that fluorescence of TagRFP675 was increased in the presence of G4s in *E. coli* but not by non-G4 sequences (Begeman *et al*., 2020) using a tightly controlled two expression vector system (**Figure 5A**). Here, we used this system for a fluorescence time-course assay to observe folding, solubility, and proteolysis after translation inhibition. Briefly, we co-expressed TagRFP675 with Seq576 RNA under heat stress. As controls, we compared the fluorescence in the presence of Seq576 both against empty vector (negative control) as well as GroEL (positive control). The expression of TagRFP675 was induced for 3 hours (**Figure 5B**), and then protein synthesis was stopped by the addition of spectinomycin, which has previously been used to halt protein translation for protein fate analyses (Barembruch and Hengge, 2007; Granot et al., 2007; Yoshitani et al., 2019).

**Figure 5.**
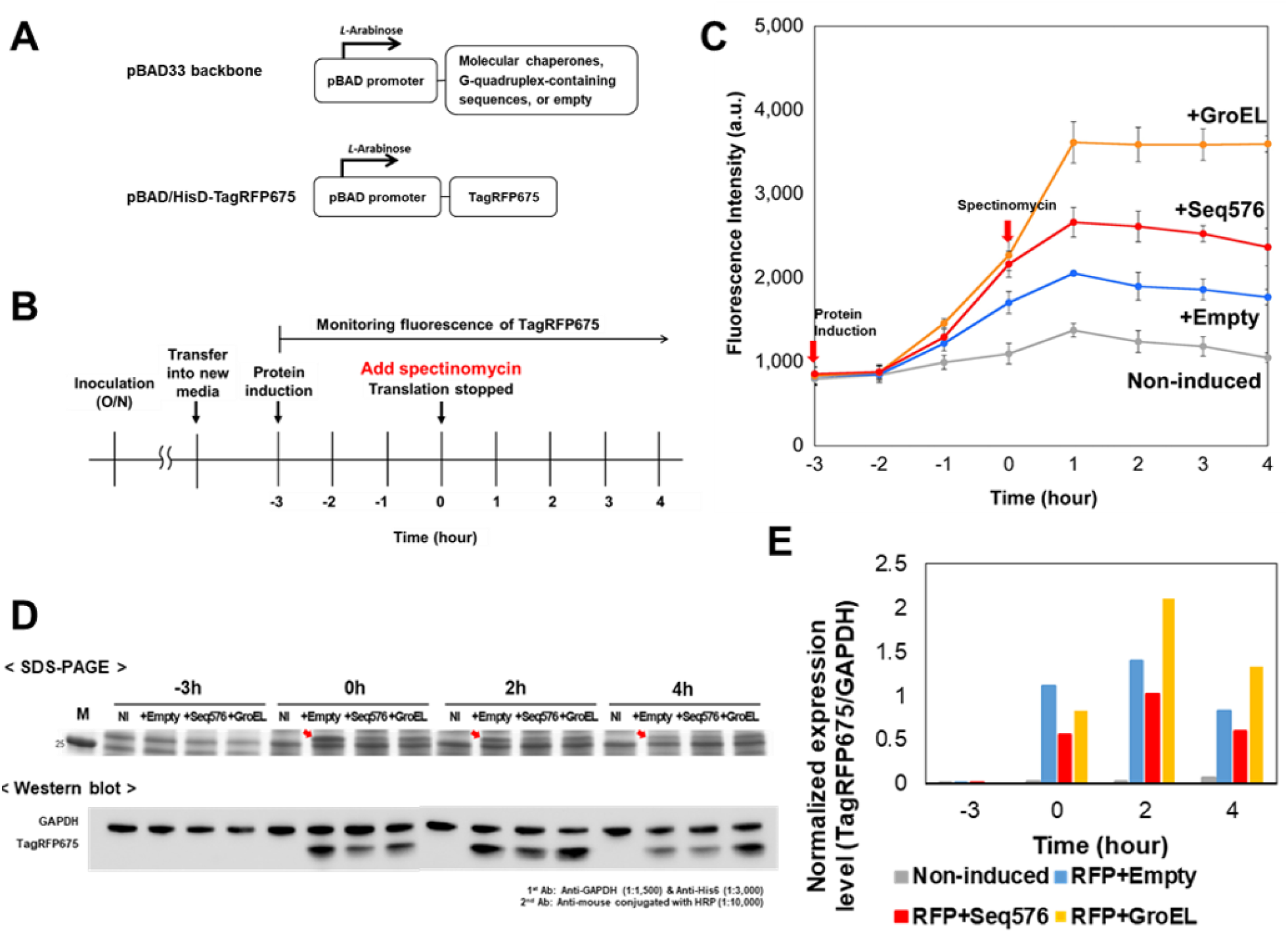
Fluorescence time-course assay of TagRFP675 in MC4100(DE3). (A) A schematic illustration of expression vectors. Both the expression of G4s and TagRFP675 are under the control of pBAD promoter, which is induced by *L*-Arabinose. (B) An experimental workflow of the fluorescence time-course assay. Protein was induced when OD_600_ reaches more than 0.7 (at time point -3h) and expressed for 3h at 42°C, and then protein translation was stopped by adding spectinomycin (at time point 0h). The fluorescence of TagRFP675 was continuously monitored after protein induction. (C) Time-course assay of TagRFP675 in the presence of G4 (Seq576) or molecular chaperone (GroEL). Protein translation was halted at time point 0h after 3h induction of protein. The experiment shown here were technical triplicates (n = 3) and performed more than 2 times. (D) SDS-PAGE (top panel) and western blot assay (bottom panel) from the cells at time points -3h, 0h, 2h, and 4h of the time-course assay. Red arrows indicate expressed TagRFP675. The concentrations of 1^st^ Anti-GAPDH, 1^st^ Anti-His6, and 2^nd^ Anti-mouse Antibodies are listed at bottom right. The full images are shown in **Figure S3**. (E) Normalization of the expression level of TagRFP675. Normalized ratio was calculated by dividing the expression level of TagRFP675 by that of GAPDH.

We observed that the fluorescence of TagRPF675 continued to increase after inhibition of translation by about a factor of two (**Figure 5C**). We confirmed that this result is not due to insufficient translation inhibition by testing varying concentrations of spectinomycin (**Figure S2A**) and other inhibitors such as kanamycin (data not shown). This behavior is instead consistent with the slow folding and maturation time of TagRFP675 (Piatkevich *et al*., 2013). Consistent with *in vitro* refolding, co-expression of the G4-containing sequence Seq576 strongly increased the fluorescence of TagRFP675 above empty vector, indicating a higher level of properly folded TagRFP675 in cells (**Figure 5C**).

To determine whether Seq576 has a role in the expression level of TagRFP675 in cells, we harvested and visualized amount of TagRFP675 present at each time point with SDS-PAGE and western blot (**Figure 5D&E**). The expression level of TagRFP675 was not significantly changed shortly after translation inhibition (0h to 2h), but it did decrease at later timepoints (4h), suggesting proteolysis was slowly decreasing the protein level. This result correlated with the slight decrease of the fluorescence at the 4h timepoint (**Figure 5C**). Curiously, the expression level of TagRFP675 was not perfectly proportional to the fluorescence. We further confirmed that there were not significant differences between pH, the translation inhibition had not killed the cells within this timeframe (**Figure S2B-D**), and the fluorescence of expressed TagRFP675 did not affect overall absorbance spectra and OD_600_ of cells for normalization (**Figure S4**). Taken together, these data suggest that not all translated TagRFP675 formed properly folded structure due to its slow folding and maturation time, and Seq576 promoted some portion of the protein population to fold.

Previous studies have reported that nucleic acid binding stimulates *E. coli* Lon protease activity, and Lon protease itself is a well-known nucleic acid-binding protein that binds a variety of nucleic acid sequences and structures (Ambro et al., 2012; Chen et al., 2008; Chung and Goldberg, 1982; Karlowicz et al., 2017; Kubik et al., 2012). As we observed degradation of TagRFP675 at later time points, the Seq576 could be potentially affecting either the activity of Lon protease or protein folding, or both. We therefore performed the same experiment in a BL21(DE3), a strain that notably lacks Lon (**Figure 6**).

**Figure 6.**
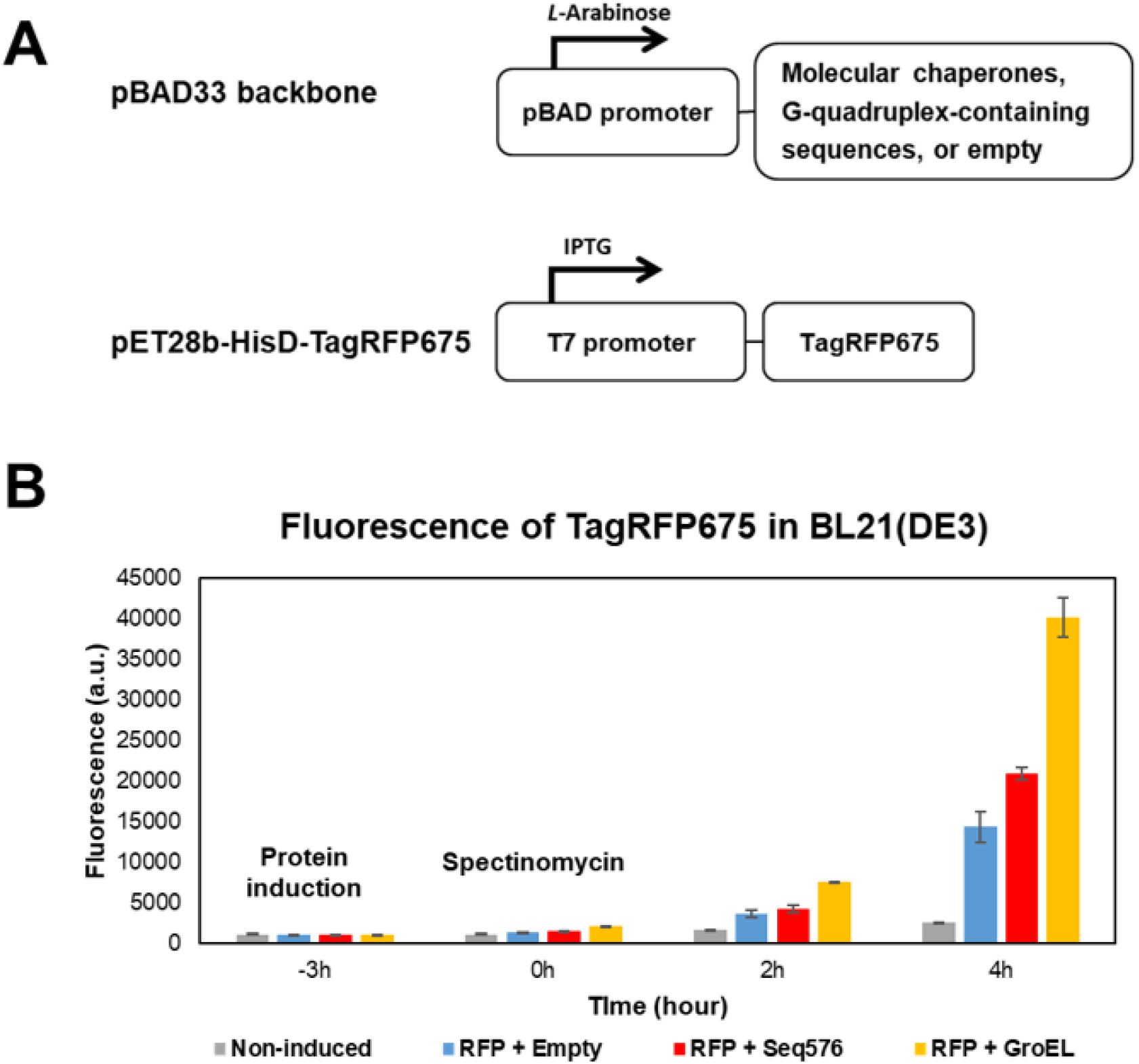
Fluorescence time-course assay of TagRFP675 in BL21(DE3). (A) A schematic illustration of expression vectors. The expression of G4s is under the control of pBAD promoter, which is induced by *L*-Arabinose. TagRFP675 expression is under the control of T7 promoter, which is induced by IPTG. (B) Time-course assay of TagRFP675 in the presence of G4s (Seq576) or molecular chaperone (GroEL). The experiment shown here were technical triplicates (n = 3) and performed more than 2 times.

In BL21(DE3), we also observed that in the presence of Seq576, the fluorescence of TagRFP675 was significantly higher than that with empty vector control, consistent with the result in MC4100(DE3). This suggests that Seq576 mainly affects folding of the protein, and not in preventing degradation by Lon. Similar to MC4100(DE3), in BL21(DE3) the fluorescence continued to increase after translation inhibition, but for a much greater timespan than in MC4100(DE3). We again confirmed that in BL21(DE) cell growth, pH, and viability between samples are similar (**Figure S5**) and GFPwt fluorescence is unaffected by Seq576 (**Figure S6**). These data further suggest that slow folding and maturation of TagRFP675 also continue in BL21(DE3), but due to reduced degradation, a larger proportion of the TagRFP675 is chaperoned and folded to the native state.

Next, we investigated whether the expression level of TagRFP675 was affected by the presence or absence of Seq576 in BL21(DE3) (**Figure 7**). Even though the difference of fluorescence between co-expressed GroEL or Seq576 and empty vector was large, the solubility and protein levels of TagRFP675 among all samples were very similar (**Figure 7A to C**). These results indicate that the primary effect of Seq576 is on enhancing proper folding, not increasing the amount of protein or its solubility. Normalizing the fluorescence by expression level confirms this observation (**Figure 7D**). Taken together with *in vitro* results, the mechanisms of G4s in assisting folding appear to enhance proper folding by helping protein intermediates to overcome kinetic traps in their protein folding landscapes.

**Figure 7.**
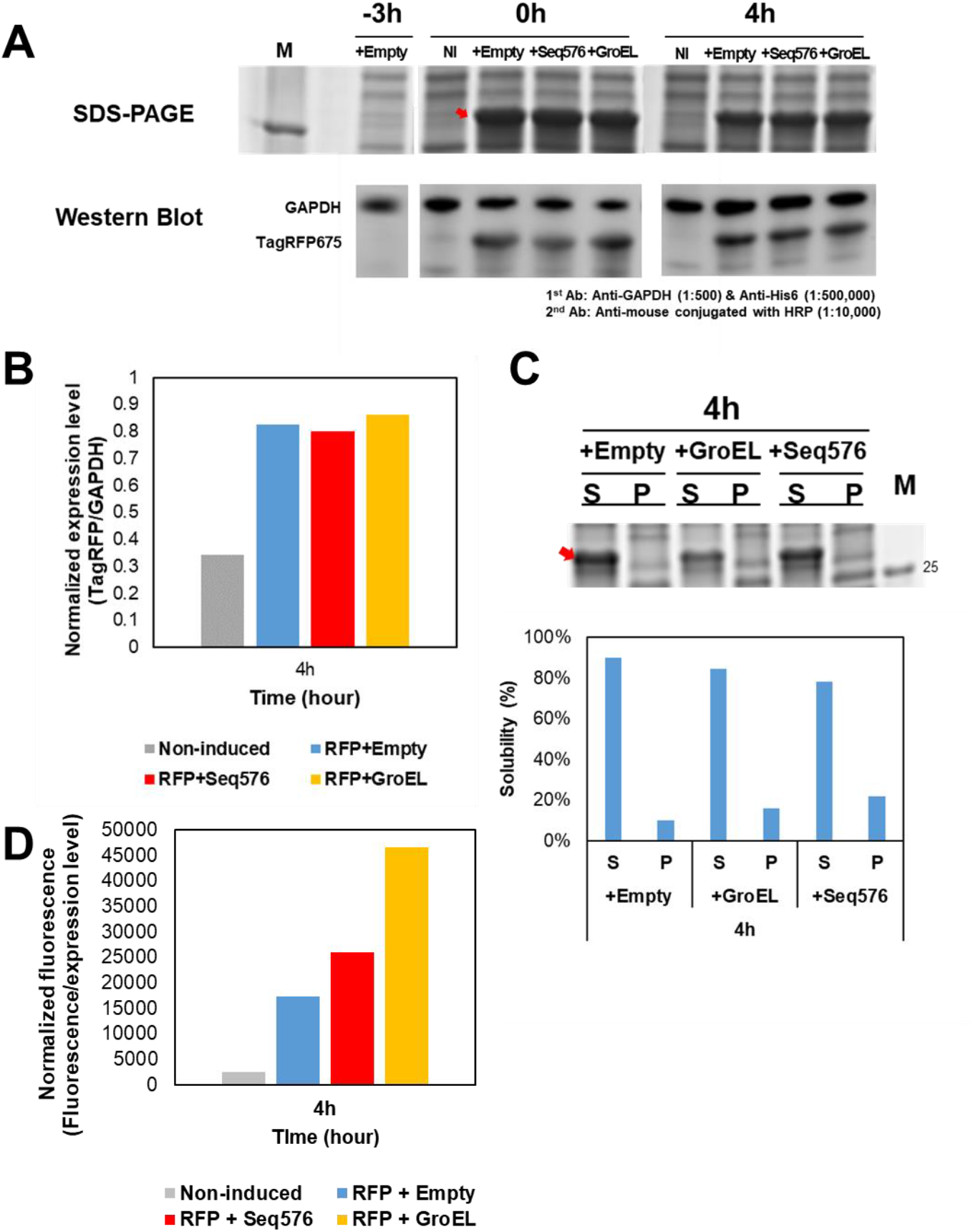
Normalization of expression level, fluorescence, and solubility of TagRFP675 from the fluorescence time-course assay in Fig 8 in BL21(DE3). (A) SDS-PAGE (top panel) and western blot assay (bottom panel) from the cells at time points -3h, 0h, and 4h of the time-course assay. A red arrow indicates expressed TagRFP675. The concentrations of 1^st^ Anti-GAPDH, 1^st^ Anti-His6, and 2^nd^ Anti-mouse Antibodies are listed at bottom right. The full images of gels are shown in **Figure S7C**. (B) Normalization of the expression level of TagRFP675 at 4h timepoint. The expression ratio was calculated via Image J. Normalized ratio was calculated by dividing the expression level of TagRFP675 by that of GAPDH. (C) Solubility of TagRFP675 at 4h timepoint. Cell lysates taken from 4h timepoints were separated into Soluble (S) and Pellet (P) fractions by centrifugation, and visualized on a SDS-PAGE gel (top). A red arrow indicates induced TagRFP675. The solubility was measured using ImageJ and shown in the bottom panel. (D) Normalization of the fluorescence of TagRFP675 at 4h timepoint. The ratio was calculated by dividing the fluorescence of induced TagRFP675 by the normalized expression level from (B). The full images of (C) and (D) are shown in **Figure S8**.

To further test whether Seq576 affects Lon activity, we added Lon protease back into BL21(DE3) cells (**Figure S7**). With Lon expression, both fluorescence and expression level of TagRFP675 in BL21(DE3) were highly decreased and were more similar to that in MC4100(DE3). Of note, we found that the increase in fluorescence by Seq576 compared to Empty vector was proportional both with and without Lon protease. This result suggests that the primary effect of Seq576 is not to prevent degradation by Lon, and is consistent with the increase in fluorescence in the presence of Seq576 not relying on an increased amount of protein.

## Discussion

In this study, we examined the effects of G4s with chaperone activity on the folding of an unstable biosensor protein, TagRFP675. *In vitro*, we unexpectedly found that the G4s accelerate protein folding to native and near-native states by rescuing kinetically trapped intermediates. In *E. coli*, we found that the G4 Seq576 improves protein folding of TagRFP675 as opposed to affecting protein solubility or proteolysis. Together, these results illuminate the potential power of G4s to modulate protein folding.

The ability for G4s to accelerate protein folding *in vitro* is somewhat surprising based on literature on protein folding rates and chaperone activity. Several ATP-independent chaperones have been observed to improve protein folding yield, such as Spy and SecB (Horowitz *et al*., 2018; Huang et al., 2021). However, in these cases the rate of folding slows down due to the affinity of partially folded intermediates for the chaperone (Horowitz *et al*., 2018; Wu et al., 2019). That said, ATP-independent “holdase” chaperones have been shown to have the capacity to potentially reshape folding landscapes (Huang *et al*., 2021; Mashaghi et al., 2013; Moayed et al., 2020). The observation that the G4 rescues kinetically trapped intermediates to both native and fluorescent non-native states suggests that the G4 is acting in a fashion that is general, but this still could have several different molecular explanations, including modification of multimeric state. These effects could come from destabilization or catalysis-like effects (**Figure 8**). Of note, it is also possible that similar effects could occur with ATP-independent chaperone proteins.

**Fig 8:**
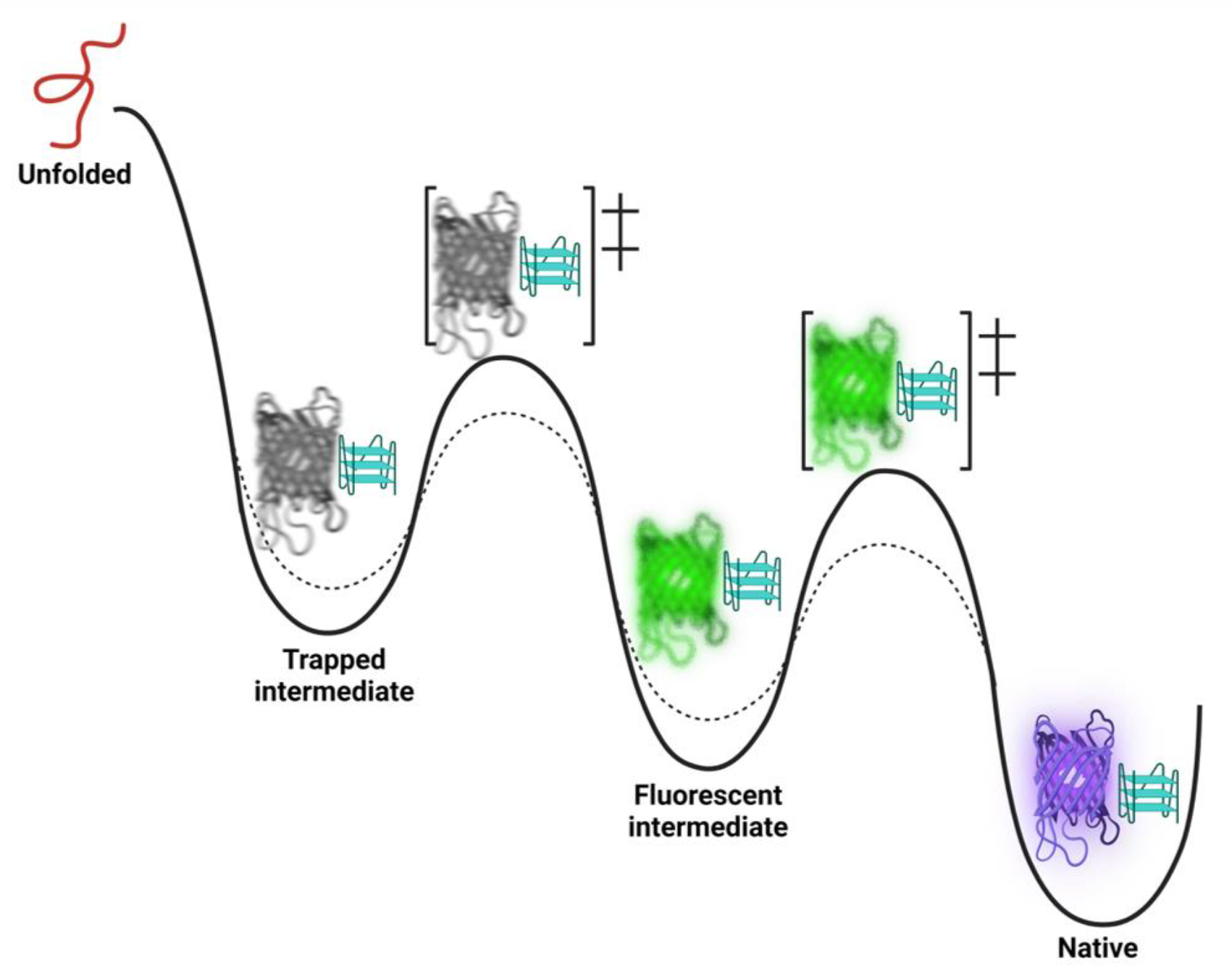
Simplified model of the effects of G4 on the folding of TagRFP675. The dashed line represents the two possible effects of the G4 to rescue protein folding. The G4 rescues protein folding either by stabilizing the transition state between the trapped intermediate, later intermediate (green), and native states (purple), or partially destabilizing the trapped intermediate state, or both. Trapped intermediate and transition state are depicted here as fuzzy protein states, while the G4 is shown in cyan.

From the *E. coli* experiments, we found that the G4s affected protein folding primarily, and not protein level or protein solubility. A common assumption made in biological experiments is that soluble protein is typically well-folded and active. However, several studies have pointed out that this is an oversimplification (Danielsson et al., 2015; Ebbinghaus et al., 2010; Nissley et al., 2022; Waztor et al., 2020), as appears to be the case for TagRFP675 when not enough chaperone capacity is available to aid its folding.

In combination with recent work on the stability of G4s under stress conditions, we can now present a potential model for the cycle of G4s in protein folding (**Figure 9**). Under stress conditions, G4s preferentially fold (Kharel et al., 2022). The protein binds to the G4 under stress, and interaction with the G4 rescues kinetically trapped intermediates. It is possible that this could cause a release of the G4, but for those proteins that stay bound, the cessation of stress causes helicase-mediated unfolding of the G4s (Kharel *et al*., 2022) and release bound proteins. This cycle could then be repeated if stress recurs. It is also possible that the G4s could still play other roles in proteostasis in addition to protein folding.

**Fig 9.**
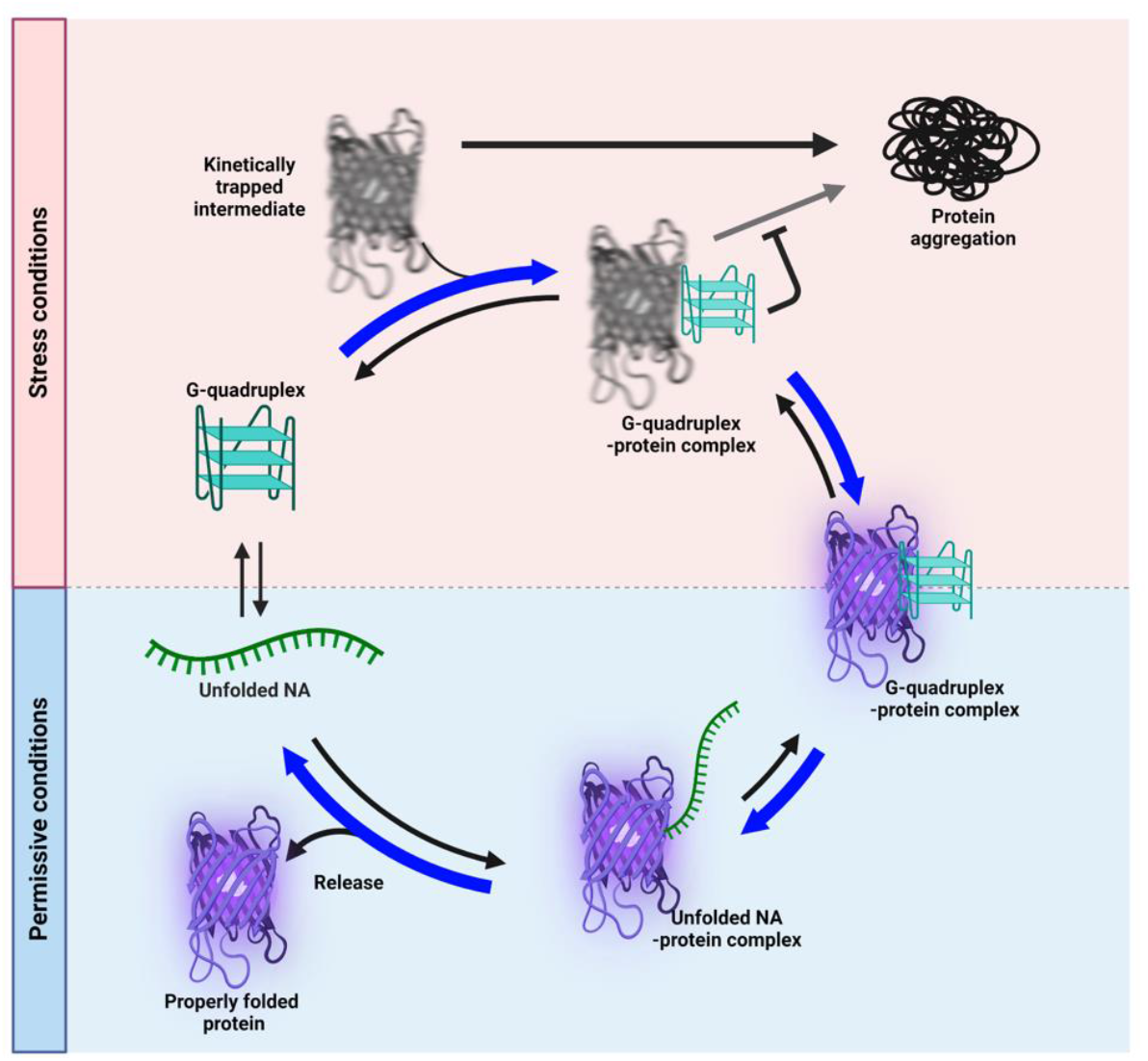
Mechanistic Folding Cycle of G4 Clients. Under stress condition, unfolded single-stranded G4-sequence-containing nucleic acid (NA) tends to form G4 sequences. Kinetically trapped or misfolded intermediates are aggregation-prone without aid of G4s. G4s bind to the intermediates, enhancing proper protein folding. Under non-stress conditions, G4s are likely to unfold, which favors release of properly folded protein from NA-protein complex.

Concluding, the role of G4s in protein folding and aggregation is still an evolving area of study, but ATP-independent chaperones such as G4s can play potentially substantial roles in dictating the folding state of proteins through rescuing trapped intermediates.

## Acknowledgements

The authors would like to thank J. Bardwell and U. Jakob for helpful discussions and F. Stull and K. Ghosh for commentary on the manuscript. This work was funded by NIH R35GM142442

## Author contributions

A.S., V.H.C., and T.L conducted experiments; A.S., T.L., and S.H. designed the experiments,, S.H. and A.S. wrote the paper; edited by all authors.

## Material and Methods

### Construction of expression vectors and nucleic acid sourcing

Strains and plasmids are listed in Table 1. pBAD/HisD-TagRFP675, pET28b-His6-TagRFP675 and pBAD18-wtGFP were used for the expression of folding biosensors. pBAD/HisD-TagRFP675 was a gift from Vladislav Verkhusha (Addgene plasmid # 44274; http://n2t.net/addgene:44274; RRID:Addgene_44274) (Piatkevich *et al*., 2013). pET28b-His6-TagRFP675 was generated from pET28b vector, using NcoI/EcoRI restriction enzymes. pBAD18-wtGFP was given by Jonathan S. Weismann’s laboratory (Wang et al., 2002). pBAD24-Lon WT was given by Igor Konieczny’s laboratory (Karlowicz *et al*., 2017). pBAD24-empty vector was generated by using EcoRI and HindIII restriction enzymes, and then blunting and ligating the cleaved vector. pBAD33 vector was used to generate expression vectors of protein folding enhancing factors, i.e., pBAD33-GroEL, pBAD33mut-Seq576, and pBAD33mut-Empty. pBAD33-GroEL was constructed using the SacI and HindIII sites and the GroEL gene was obtained by PCR using *E. coli* genome. pBAD33mut-empty vector was generated from pBAD33 vector by changing MCS into SacI and SpeI, using MluI and BglII restriction enzymes. The genes of Seq576 was synthesized (Genscript) and inserted into pBAD33mut vector using the SacI and SpeI sites. DNAs for *in vitro* refolding assays were synthesized (IDT) and annealed before use.

**Table 1.** Bacterial strains and plasmids.

| Strains |  | Genotype |  |  |  | References |  |  |
| --- | --- | --- | --- | --- | --- | --- | --- | --- |
| MC4100(DE3)<br>BL21(DE3) | | $\Delta(argF-lac)$ *U169 araD139 rpsLS50 reLAI deoCI ptsF25 rpsR flbB301 (DE3)*<br>*fhuA2 [lon] ompT gal ($\lambda$ DE3) [dcm] ΔhsdS* | | | | (Casadaban, 1976)<br>(Studier and Moffatt, 1986) | | |
| Strain number | Strains | Plasmid1 | Relevant Characteristics | Plasmid2 | Relevant Characteristics | Plasmid3 | Relevant Characteristics | References |
| AS181 | MC4100(DE3) | pBAD/HisD-TagRFP675 | ApR | pBAD33mut-Empty | CmR |  |  | (Begeman *et al.*, 2020) |
| AS183 | MC4100(DE3) | pBAD/HisD-TagRFP675 | ApR | pBAD33-GroEL | CmR |  |  | (Begeman *et al.*, 2020) |
| AS197 | MC4100(DE3) | pBAD/HisD-TagRFP675 | ApR | pBAD33mut-Seq576 | CmR |  |  | (Begeman *et al.*, 2020) |
| AS491 | MC4100(DE3) | pBAD19-wtGFP | ApR | pBAD33mut-Empty | CmR |  |  | This study |
| AS497 | MC4100(DE3) | pBAD19-wtGFP | ApR | pBAD33-GroEL | CmR |  |  | This study |
| AS493 | MC4100(DE3) | pBAD19-wtGFP | ApR | pBAD33mut-Seq576 | CmR |  |  | This study |
| AS428 | BL21(DE3) | pET28b-His6-TagRFP675 | KanR | pBAD33mut-Empty | CmR | pBAD24-Empty | ApR | This study |
| AS563 | BL21(DE3) | pET28b-His6-TagRFP675 | KanR | pBAD33-GroEL | CmR | pBAD24-Empty | ApR | This study |
| AS430 | BL21(DE3) | pET28b-His6-TagRFP675 | KanR | pBAD33mut-Seq576 | CmR | pBAD24-Empty | ApR | This study |
| AS385 | BL21(DE3) | pET28b-His6-TagRFP675 | KanR | pBAD33mut-Empty | CmR | pBAD24-Lon WT | ApR | This study |
| AS565 | BL21(DE3) | pET28b-His6-TagRFP675 | KanR | pBAD33-GroEL | CmR | pBAD24-Lon WT | ApR | This study |
| AS389 | BL21(DE3) | pET28b-His6-TagRFP675 | KanR | pBAD33mut-Seq576 | CmR | pBAD24-Lon WT | ApR | This study |
| AS524 | BL21(DE3) | pBAD19-wtGFP | ApR | pBAD33mut-Empty | CmR |  |  | This study |
| AS527 | BL21(DE3) | pBAD19-wtGFP | ApR | pBAD33-GroEL | CmR |  |  | This study |
| AS526 | BL21(DE3) | pBAD19-wtGFP | ApR | pBAD33mut-Seq576 | CmR |  |  | This study |

### Protein expression and purification

TagRFP675 was expressed in BL21(DE3) bacterial host cells with plasmids pBAD/HisD-TagRFP675 and co-expressing pBAD33-GroEL. Cells were grown in Luria-Bertani (LB) medium with ampicillin and chloramphenicol for 24 hours at 37°C while shaking at 180 rpm. Afterwards, a starter culture was transferred to fresh LB medium containing the previous antibiotics at 37°C while shaking at 250 rpm, until an OD_600_ of 0.7 was achieved. Cells were induced with 0.2% *L*-arabinose and allowed to incubate at 250 rpm for 45 minutes or until spun down aliquots had a distinct red/purple pellet. Cells were collected by centrifugation at 6,000 rpm for 20 minutes. The supernatant was discarded while the pellet was resuspended in 50 mM sodium phosphate, 300 mM NaCl, 10 mM imidazole, 10mM PMSF, and autoclaved DI water. Cells were then lysed by sonication on ice. The lysate was centrifuged for 30 min at 15,000 g and the supernatant was subsequently filtered using 0.45 μm micron syringe filters. The recombinant protein was purified using Bio-Scale Mini Nuvia IMAC Ni-Charged column. The protein was eluted through a gradient elution with an elution buffer made of 50 mM sodium phosphate, 300 mM NaCl, and 300 mM imidazole. The loading buffer used consisted of 50 mM sodium phosphate, 300 mM NaCl, and 10 mM imidazole. Purification was followed by dialysis overnight in 10 mM sodium phosphate, pH 7.5, and these were then concentrated using a Amicon Ultra-15 Centrifugal Filter Unit, being spun at intervals of 20 minutes at 20,000 rpm. After the 20 minutes intervals the sample was pipetted to reach a homogenous solution, and this was done until the desired concentration and volume was achieved. The protein was then flash frozen with 15% glycerol for storage. Purity was assessed through an SDS-PAGE at greater than 95% purity. **Figure S9** shows the gel portraying the purified protein sample.

### Protein Denaturation and Fluorescence Recovery Assay

The fluorescence intensity of renaturing TagRFP675 was measured with a fluorimeter in the presence and absence of G4 containing nucleic acids. TagRFP675 was denatured chemically with 6 M guanidine-HCl, 40 mM HEPES, pH 7.5 (KOH) for approximately 11 hours at 23°C in tear drop vials at 20 µM protein. Denatured protein was then diluted into 10mM potassium phosphate, pH 7.5, in a clear cuvette and with constant stirring at 23°C. Fluorescence was measured by the fluorimeter set at an emission wavelength of 675 nm, an excitation wavelength of 598 nm, high detector voltage (800 V), and 10 nm slits on excitation and emission. When measuring the fluorescence recovery in the presence of nucleic acid, the nucleic acid was first added to the 10 mM potassium buffer in the cuvette, followed by the dilution of the denatured protein into the cuvette after 1 minute of starting the readings in the fluorimeter. All samples and conditions were run in triplicates and graphed in an intensity vs time plot.

### Fluorescence Recovery: temperature

Measurements were recorded at 4°C and 40°C. The same procedure was followed as the fluorescence recovery assay in the presence of nucleic acids. Seq576 was present in the potassium buffer prior to the dilution of the denatured TagRFP675. The concentration ratio of TagRFP65 to Seq576 was 1:2. For all temperatures, the buffer was allowed to rest in the cuvette holder for approximately 2 minutes before diluting protein in buffer.

### Fluorescence Recovery: delayed addition of protein

Seq576 was added to the sample in the cuvette before and after the dilution of the denatured TagRFP675. Seq576 was added before dilution (0 seconds), or at later time points indicated in figure after TagRFP675 was diluted in the native buffer. The fluorescence recovery was taken at 23°C.

### Fluorescence assay in Escherichia coli

Each resulting expression vector of protein folding enhancing factors and pBAD/HisD-TagRFP675 and pET28b-His6-TagRFP675 were co-transformed into the *E. coli* strain MC4100(DE3) or BL21(DE3) by heat shock. To perform time-course fluorescence assay, the same expression vector set mentioned above were used (**Figure 5A&6A**). Briefly, each transformant was inoculated into 3 ml LB media with appropriate antibiotics and incubated at 37°C overnight with vigorous shaking. Next day, 200 μl of each culture was transferred into fresh 3 ml LB with the same concentration of antibiotics used above. When the OD_600_ reaches > 0.7, 0.1% *L*-Arabinose was added to induce protein and RNA expression in MC4100(DE3), and 0.1% *L*-Arabinose and 100 nM IPTG were used for the expression in BL21(DE3) (referred as -3h). After 3-hour induction (referred as 0h), translation was stopped by adding excess amounts of Spectinomycin (0.5 mg/ml). As high concentrations of spectinomycin ranged from 250 µg/ml to 1mg/ml were used from previous studies (Barembruch and Hengge, 2007; Granot *et al*., 2007; Yoshitani *et al*., 2019), we first performed time-course assay at the various concentrations of spectinomycin to establish an optimum concentration of spectinomycin for our assay (**Figure S2A**). The fluorescence of TagRFP675 with 0.1 mg/ml of spectinomycin continuously increased, while more than 0.2 mg/ml of spectinomycin halted the increase in fluorescence. Thus, 0.5 mg/ml was finally chosen for our assay to monitor the degradation of protein in bacteria. Fluorescence was measured by microplate reader (Infinite M200 Pro, Tecan) using a black 96-well plate (Corning Black NBS 3991). Fluorescence emissions of TagRFP675 at 675 nm was measured upon excitation at 598 nm.

### SDS-PAGE and Western blot

Each sample from the time-course fluorescence assay was harvested at certain time points by centrifugation at 10,000 *g* for 2 min at 4°C. Cells were lysed with EZLys™ Bacterial Protein Extraction Reagent (BioVision) by following the manufacture’s method. Lysed cells were determined by dividing the OD_600_ by three to normalize the number of cells per ml for all samples. To examine the solubility of TagRFP675, the lysates were separated into soluble and pellet fractions by centrifugation at 16,000 xg for 15 min at 4°C. The samples were denatured with the same volume of SDS loading buffer at 98°C for more than 20 min and visualized on an SDS-PAGE gel. To perform western blot, each gel was transferred to 0.2 µm PVDF membrane using Trans-Blot Turbo Transfer system (Bio-Rad). To detect TagRFP675, blots were incubated with 6x-His Tag Monoclonal Antibody (Thermo Fisher Scientific). As a control, GAPDH Loading Control Monoclonal Antibody (GA1R) (Thermo Fisher Scientific) was obtained. Rabbit anti Mouse IgG (H+L) Secondary Antibody, HRP (Invitrogen) was used for visualization of bands. The dilution ratio of three antibodies for the samples in MC4100(DE3) and BL21(DE3) are listed in the figures. The protein bands were then visualized by Clarity Max™ Western ECL Substrate (Bio-Rad) using Imaging System Chemidoc (Bio-Rad). The intensity of each band was quantified by ImageJ software (n=3) (Schneider et al., 2012), and the expression level of TagRFP675 was calculated based on the expression level of GAPDH (TagRFP675/GAPDH).

