## Supplement for "G-quadruplexes rescuing protein folding"

### Supplemental Results

To verify whether the fluorescence of induced TagRFP675 has any interfering effect on $OD_{600}$ of cells, we examined absorbance spectra of samples, and compared $OD_{600}$ with addition of purified TagRFP675 (**Figure S4**). As shown in **Figure S4A,** there are no significant differences between non-induced cells and induced cells, and the shape of graph was the same with that of standard *E. coli*. We further confirmed that TagRFP675 expression would not affect $OD_{600}$ quantification by adding purified TagRFP675 to cells without a fluorescent protein to a similar fluorescence level as cells expressing TagRFP675, and then remeasuring absorbance. This test found that the cells with added purified TagRFP675 had the same $OD_{600}$ measurement both before and after TagRFP675 protein addition (**Figure S4 B&C**).

Because we previously showed nucleic acids, including G4s, had no effect on enhancing protein folding of wild-type GFP (GFPwt) (Begeman et al., 2020), we repeated this time-course experiment with GFPwt as negative control (**Figure S6**). However, due to the slow folding and maturation time, we were not able to observe significant fluorescence from a 3 h induction in fluorescence time-course assay (**Figure S6B**). Thus, we used the plate fluorescence assay based on our previous methods (Begeman *et al.*, 2020). As previously shown, G4s do not enhance the fluorescence of GFPwt in plate fluorescence assay in MC4100(DE3) (**Figure S6C**), probably due to its highly acidic nature (pI=5.67) compared to TagRFP675 (pI=8.53). The same trends were observed in BL21(DE3), showing that the inability for Seq576 to act upon GFPwt was not strain dependent.

Due to the deficiency of major proteases, we chose BL21(DE3) for our time-course assay. To assess the effect of Lon in particular, we added Lon protease back in BL21(DE3) cells, by using triple expression vector system (**Fig S7A**) to determine if our findings in BL21(DE3) are primarily Lon-mediated. With Lon expression, both fluorescence and expression level of TagRFP675 were highly decreased, compared to empty vector control (**Fig S7B**). Interestingly, the overall fluorescence intensity of TagRFP675 was similar to that in MC4100(DE3) with the addition of Lon protease. This further supports that the decrease in fluorescence and expression of TagRFP675 in MC4100(DE3) were mainly due to proteolytic degradation. We also found that the increase in fluorescence by Seq576 was proportional in the presence and absence of Lon protease, suggesting that the G4 did not affect Lon activity. These results further support that the primary role of Seq576 in enhancing fluorescence is assisting proper folding and appears to be relatively independent of Lon.

### Supplemental Figure Legends

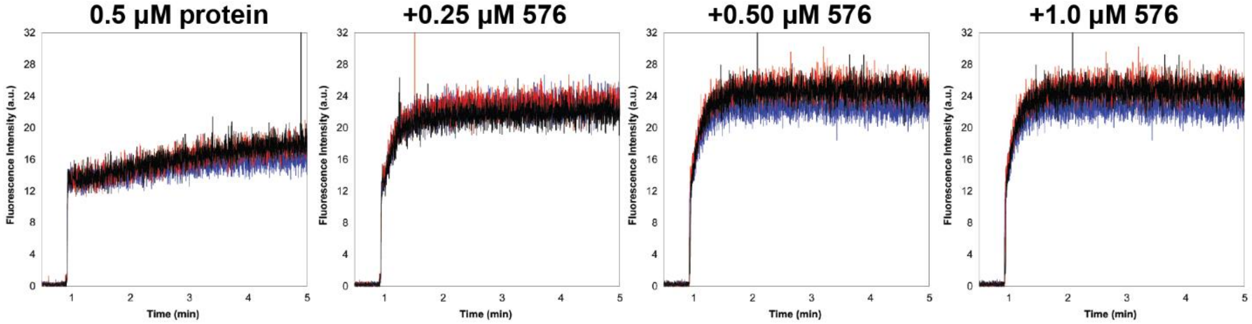


**Fig S1: Protein Folding Dependence on G4 concentration.** TagRFP675 was diluted from denaturing buffer (6 M Gu-HCl) to 0.5 μM protein into refolding buffer in the presence of different concentrations of Seq576 (at 0:1, 0.5:1, 1:1, and 2:1 ratios).

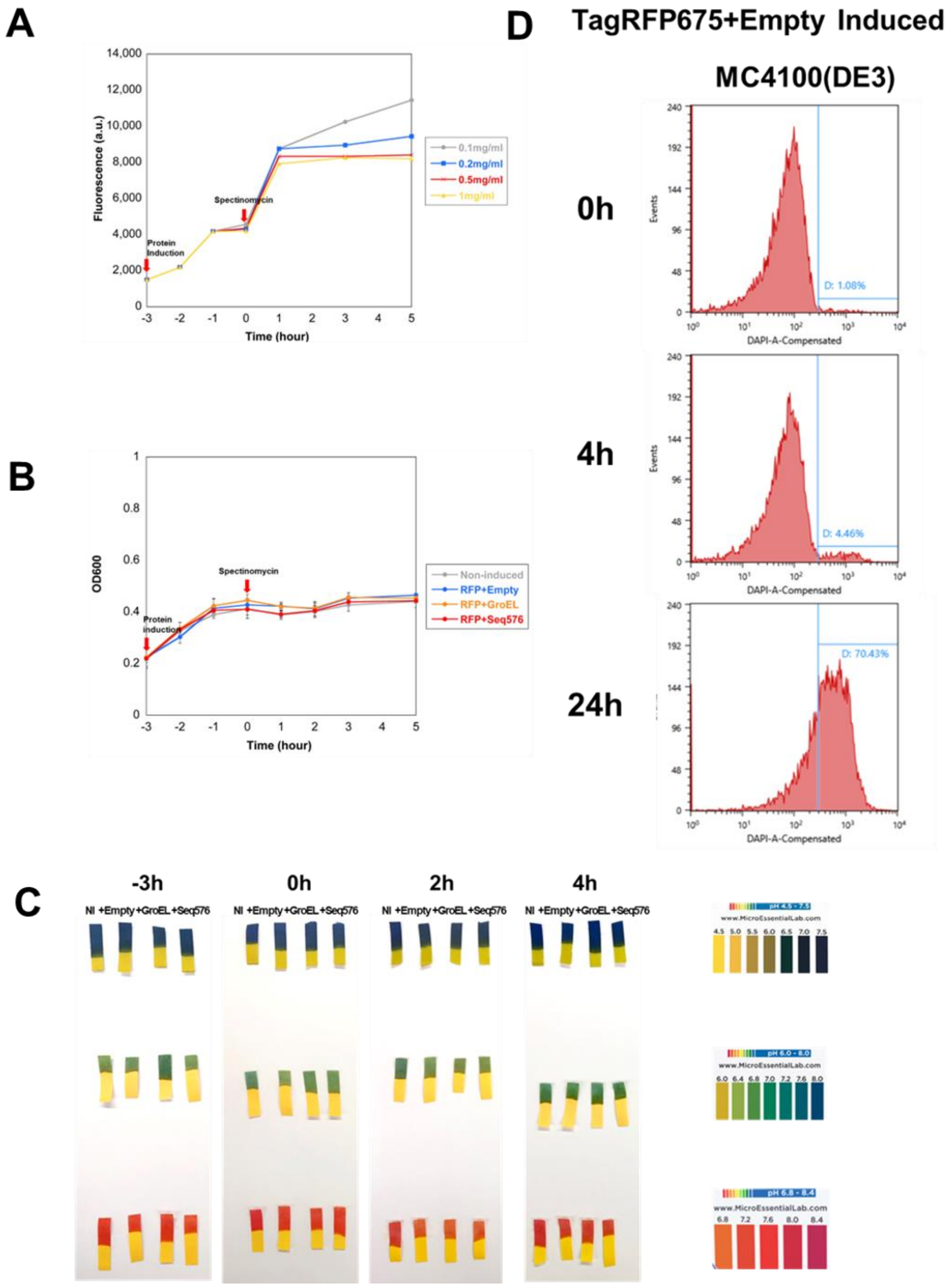


**Figure S2. Fluorescence time-course assay performed in MC4100(DE3) did not have significant effect on cell growth and viability.** (A) Time-course assay at various concentrations of spectinomycin. To check effective concentration of spectinomycin for halting protein translation, various concentrations, 0.1, 0.2, 0.5, and 1.0 mg/ml, of spectinomycin were used. 0.5 mg/ml was chosen for optimum concentration in translation inhibition. (B) Measurement of $OD_{600}$ of cells. $OD_{600}$ was measured while performing the time-course assay. (C) Measurement of pH of cell cultures. pH was measured at -3h, 0h, 2h, and 4h time points using 3 pH papers to measure various pH ranges. (D) Testing cell death by flow cytometry. DAPI (x-axis) indicates cell death. Samples from 0h, 4h and 24h time points were taken from the time-course assay.

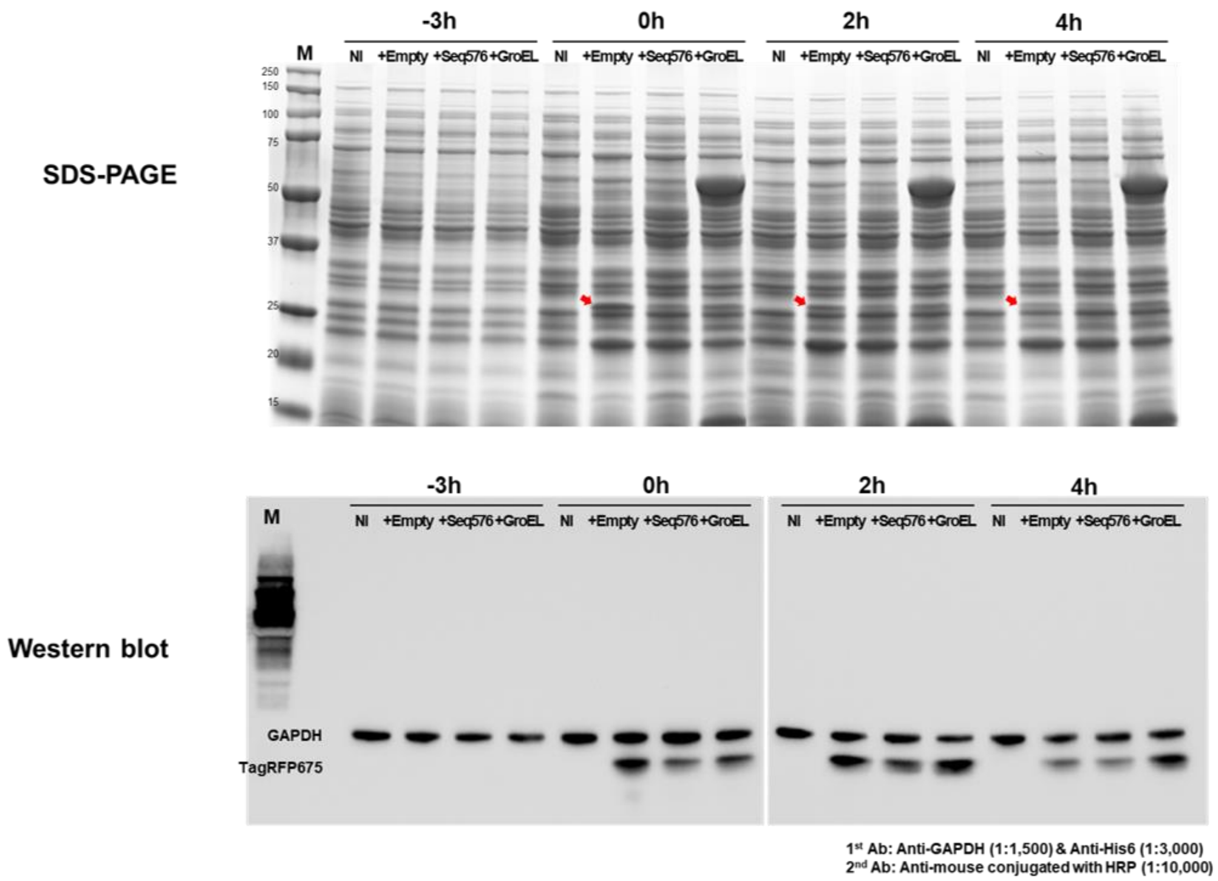


**Figure S3. Full images of SDS-PAGE (top panel) and western blot assay (bottom panel) from the cells at time points -3h, 0h, 2h, and 4h of the time-course assay.** Red arrows indicate expressed TagRFP675. The concentrations of 1st Anti-GAPDH, 1st Anti-His6, and 2nd Anti-mouse antibodies used for western blot are listed at bottom right. The cropped images were used in **Figure 5D**.

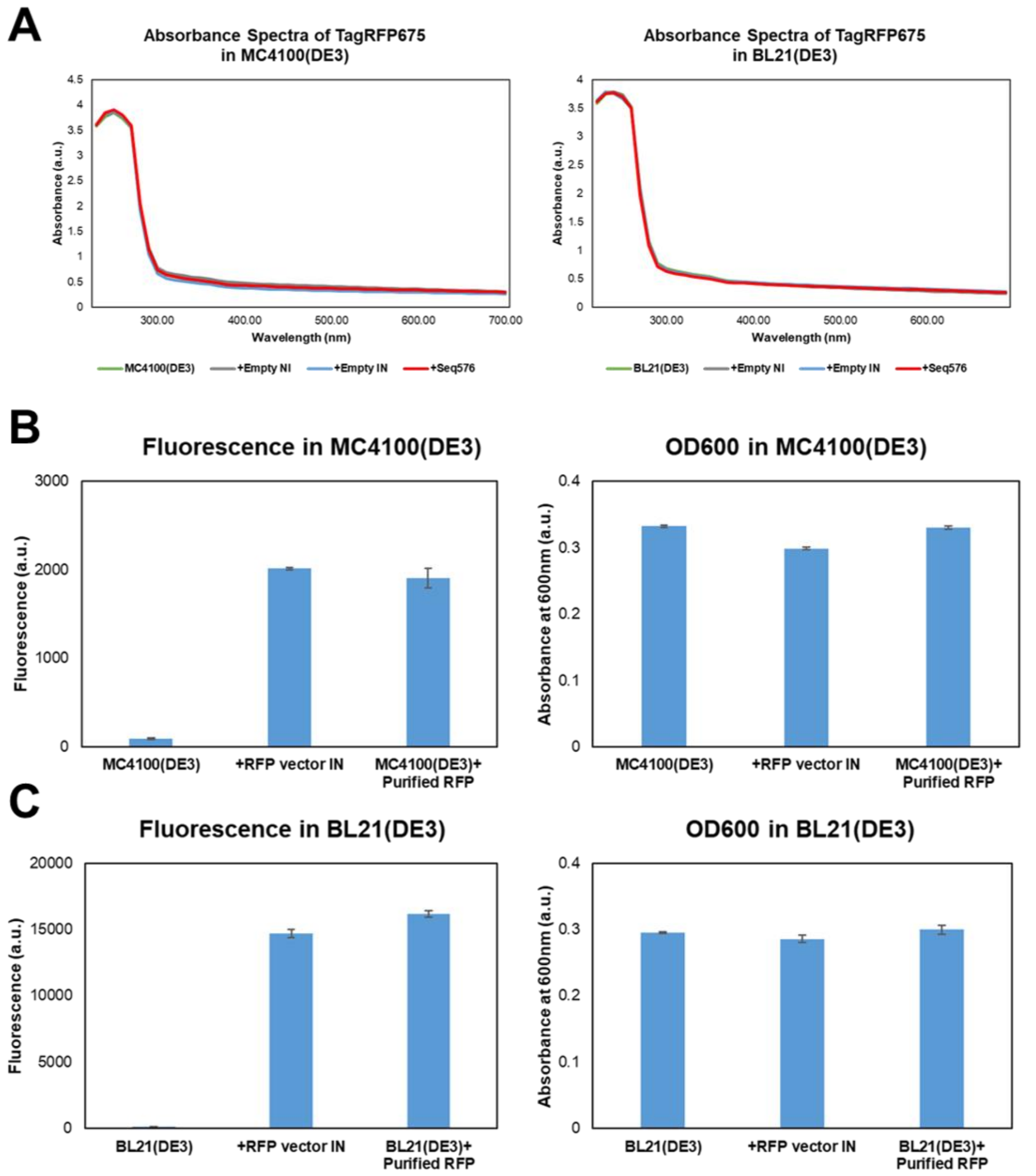


**Figure S4. Fluorescence of TagRFP675 has no significant effect on absorbance spectra and $OD_{600}$ of BL21(DE3).** (A) Absorbance spectra of TagRFP675 in the 230 nm to 1,000 nm range in MC4100(DE3) (left) and BL21(DE3) (right) were measured and compared with that of non-induced mock cells. Comparison of $OD_{600}$ between non-induced mock cells and the cells after addition of purified TagRFP675 in MC4100(DE3) (B) and BL21(DE3) (C). Both MC4100(DE3) and BL21(DE3) do not have a TagRFP675 expression vector. TagRFP675-induced cells (+RFP vector IN) were shown together to compare the fluorescence and $OD_{600}$.

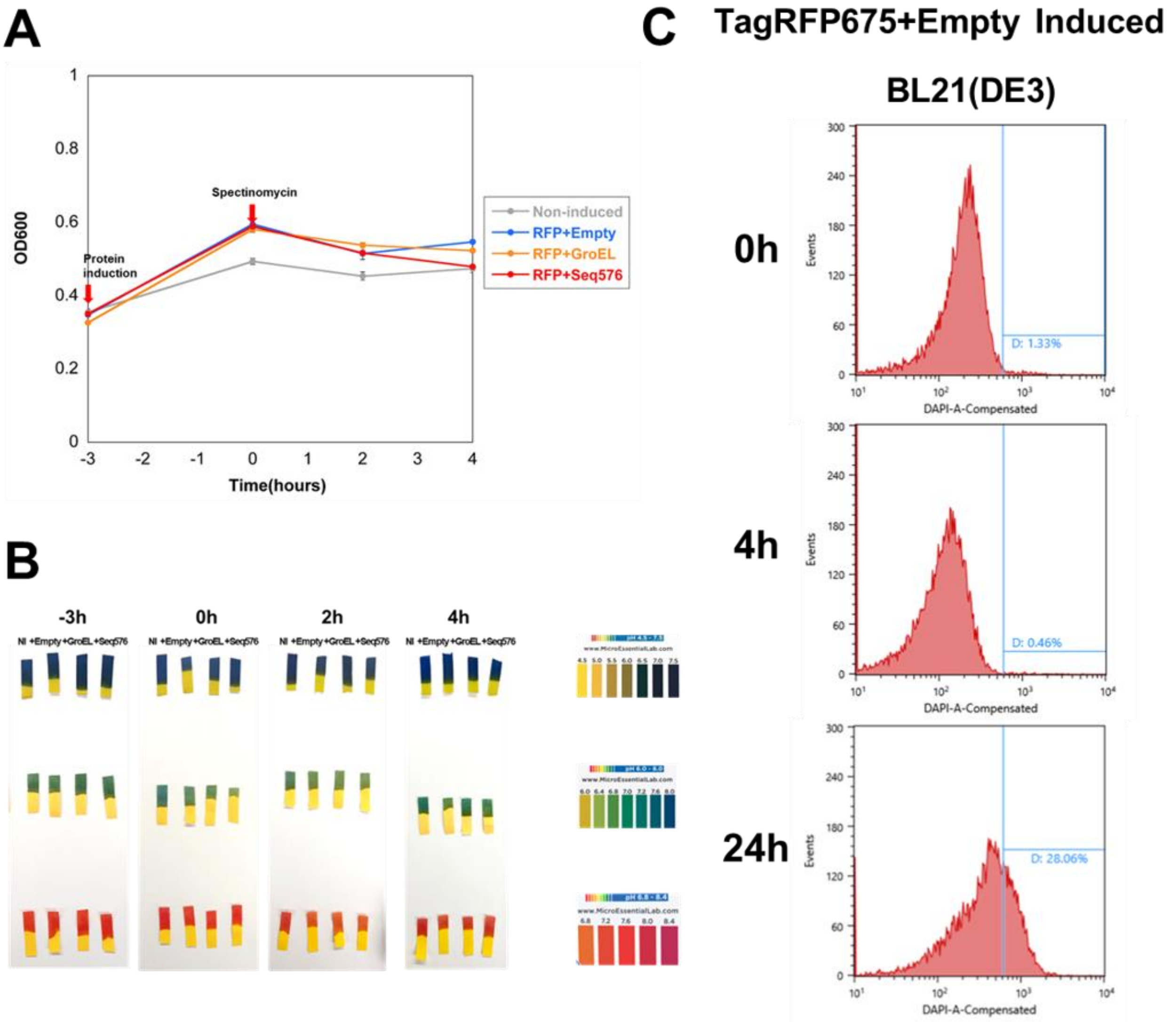


**Figure S5. Fluorescence time-course assay performed in BL21(DE3) also did not have significant effect on cell growth and viability.** (A) Measurement of $OD_{600}$ of cells. $OD_{600}$ was measured while performing the time-course assay. (B) Measurement of pH of cell cultures. pH was measured at -3h, 0h, 2h, and 4h time points using 3 pH papers to measure various pH ranges. (D) Testing cell death by flow cytometry. DAPI (x-axis) indicates cell death. Samples from 0h, 4h and 24h time points were taken from the time-course assay

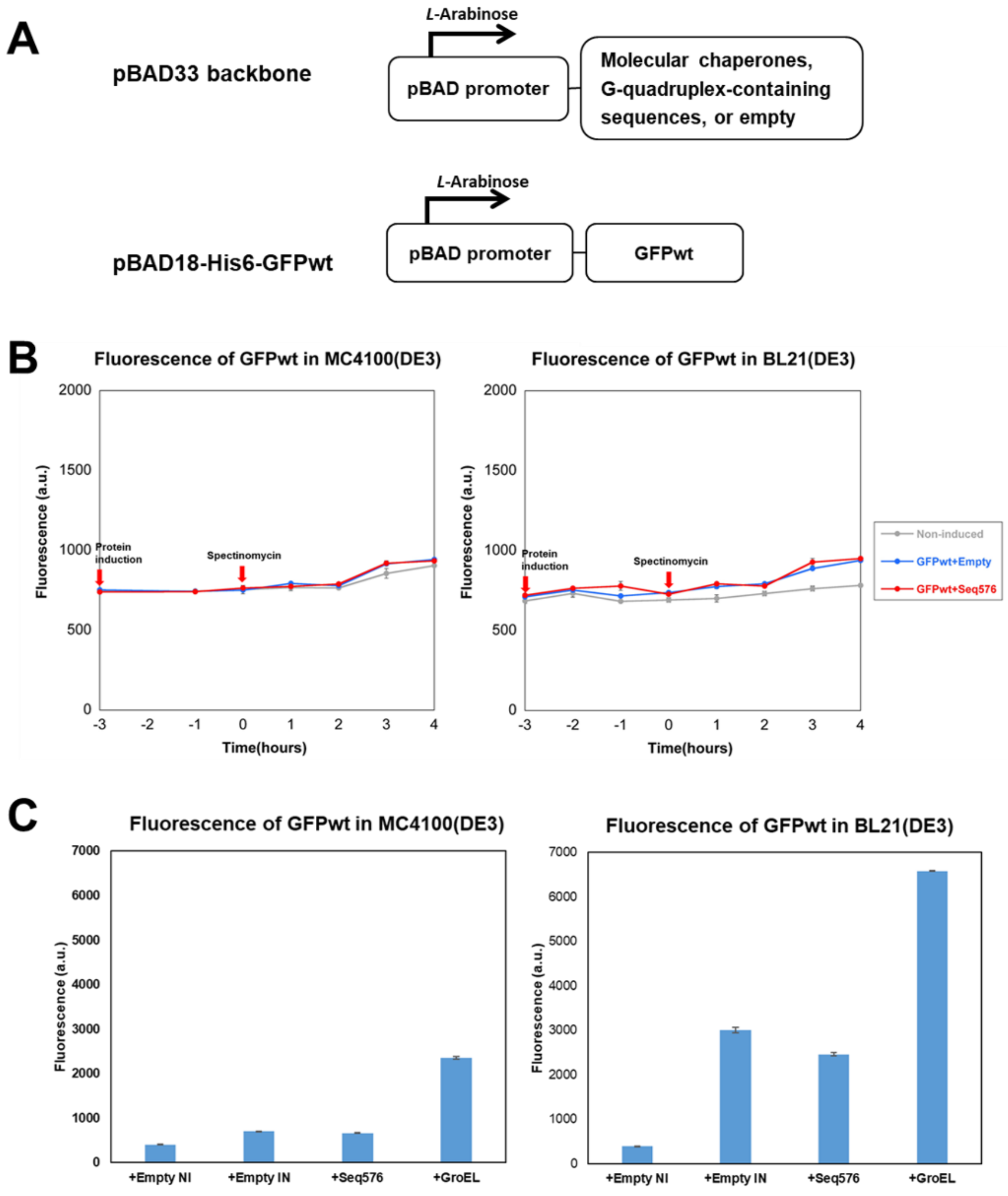


**Figure S6. G4s do not rescue the folding of GFPwt in both MC4100(DE3) and BL21(DE3)**. (A) A schematic illustration of expression vectors. Both the expression of G4s and GFPwt are under the control of pBAD promoter, which is induced by *L*-Arabinose. (B) Fluorescence time-course assay of GFPwt in MC4100(DE3) (left) and BL21(DE3) (right). (C) Fluorescence assay using LB plates in MC4100(DE) (left) and BL21(DE3) (right). GFPwt was co-expressed with empty (negative control), Seq576 (G4s), and GroEL on LB plates at 37℃ overnight. The fluorescence of each sample was measured in triplicates (n = 3).

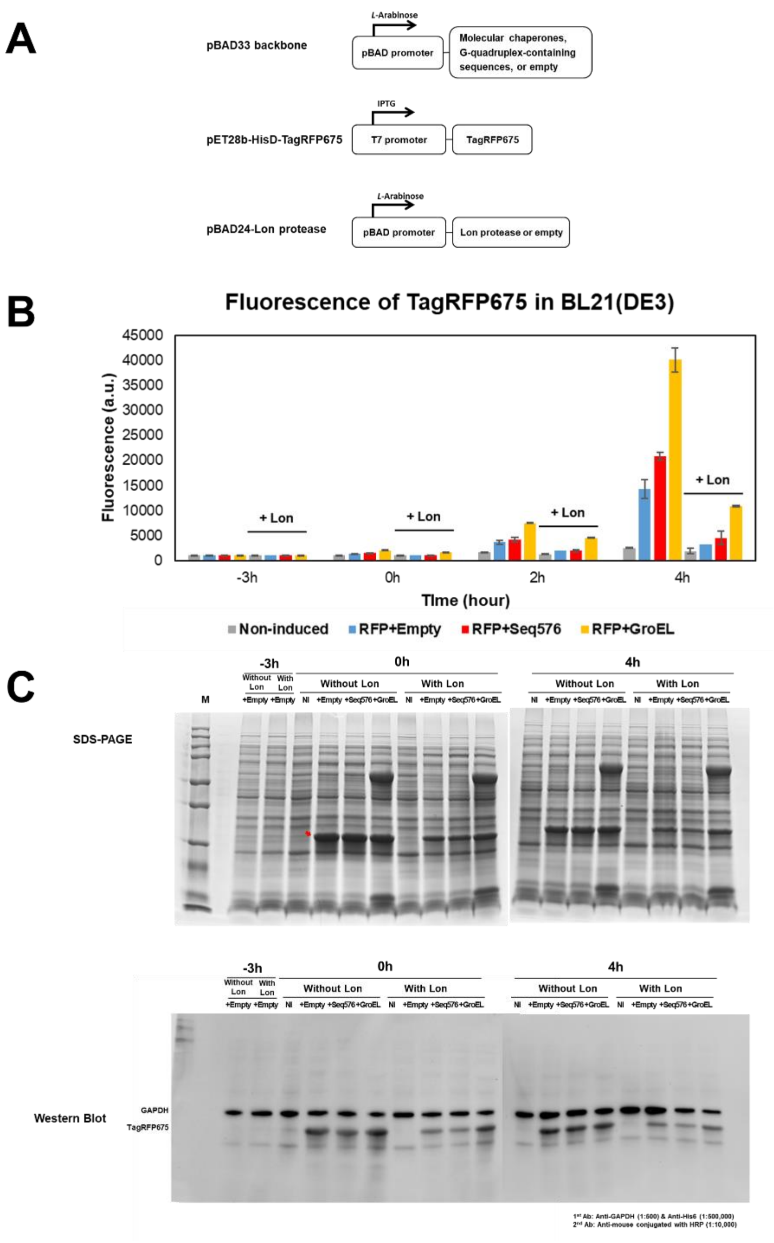


**Figure S7. Lon protease addition strongly decreases the fluorescence of TagRFP675.** (A) A schematic illustration of expression vectors. Both of the expression of G4s and Lon protease are under the control of pBAD promoter, which is induced by *L*-Arabinose. TagRFP675 was cloned into pET28b backbone, which is under the control of T7 promoter, induced by the addition of IPTG. (B) Fluorescence time-course assay of TagRFP675 in the absence (pBAD24-Empty) or presence (pBAD24-Lon) of Lon protease addition. The fluorescence of TagRFP675 was measured at timepoints -3h, 0h, 2h, and 4h. The assay shown here were technical triplicate (n = 3). (C) Full images of SDS-PAGE (top panel) and western blot assay (bottom panel) from the cells at time points -3h, 0h, and 4h of the time-course assay. A red arrow indicates expressed TagRFP675. The concentrations of 1st Anti-GAPDH, 1st Anti-His6, and 2nd Anti-mouse antibodies used for western blot are listed at bottom right. The cropped images were used in **Figure 7A**.

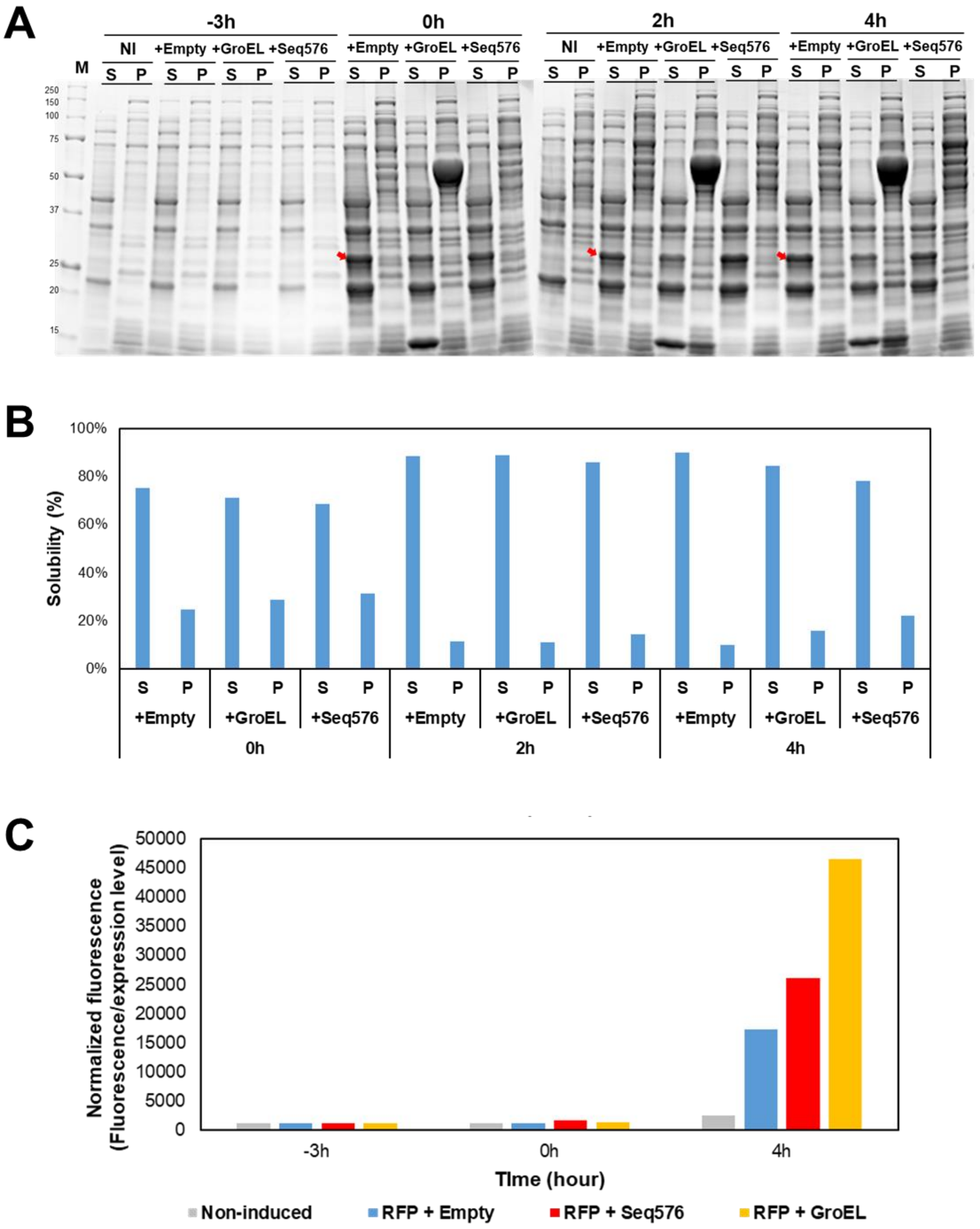


**Figure S8. Normalization of expression level, fluorescence, and solubility of TagRFP675 from the fluorescence time-course assay in BL21(DE3).** (A) Solubility of TagRFP675 at -3h, 0h, 2h, and 4h timepoints. Cell lysates were separated into Soluble (S) and Pellet (P) fractions by centrifugation and visualized on SDS-PAGE gels. Red arrows indicate induced TagRFP675. (B) The solubility was measured using ImageJ by three different people, and the average shown. (C) Normalization of the fluorescence of TagRFP675 at -3h, 0h, and 4h timepoints. The ratio was calculated by dividing the fluorescence of induced TagRFP675 by the normalized expression level. The cropped images were used in **Fig 7C&D**.

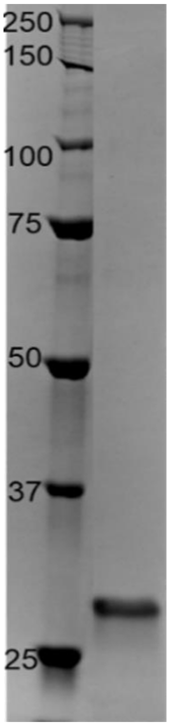


**Figure S9 Purification of TagRFP675.** The purification of TagRFP675 was analyzed through an SDS-PAGE gel after the use of native column chromatography for the protein purification. The gel shows the standard ladder used and the band corresponds to TagRFP675, with a molecular weight of 26.4kDa.

### Supplemental Material and Methods

#### *Fluorescence Recovery: concentration ratios*

Seq576 was added to the potassium phosphate buffer in the cuvette at a concentration of 0μM, 1μM, 0.5μM, and 0.25μM, resulting in concentration ratios of TagRFP675 to Seq576 of 1:0, 1:2, 1:1, and 2:1 respectively. The fluorescence recovery was recorded at 23℃.

#### *Fluorescence assay in Escherichia coli*

Each resulting expression vector of protein folding enhancing factors and pBAD/HisD-TagRFP675, pET28b-His6-TagRFP675, or pBAD33-wtGFP were co-transformed into the *E. coli* strain MC4100(DE3) or BL21(DE3) by heat shock. For the fluorescence assay in MC4100(DE3) using LB Agar plates, each transformant was spread on an LB plate containing

0.2% *L*-Arabinose, ampicillin (200 μg/ml), and chloramphenicol (50 μg/ml) at 42℃ overnight. For the assay in BL21(DE3), an LB plate containing 1μM IPTG, 0.2% *L*-Arabinose, kanamycin (50 μg/ml), and chloramphenicol (50 μg/ml) was used. To confirm whether promoters are tightly controlled, non-induced LB plates containing the same amount of antibiotics were used. The next day, cells were scraped in 170 mM NaCl and harvested by centrifugation at 10,000 *g* for 2 min at 4℃. Cells were resuspended in 170 mM NaCl and diluted to 0.1 $OD_{600}$ for further fluorescence assays. Fluorescence was measured following the same methods described in Material and Methods. Fluorescence emissions of TagRFP675 (at 675 nm) and wtGFP (at 395 nm) were measured upon excitation at 598 nm and 509 nm, respectively. Time-course assay was performed under the same set described in Material and Methods. To co-express Lon protease, three vectors shown in **Figure S7A** were co-transformed into BL21(DE3) and time-course assay were performed under the same condition above except using LB media containing, kanamycin (50 μg/ml), chloramphenicol (50 μg/ml), and ampicillin (200 μg/ml). All experiments were performed in triplicate.

#### *Escherichia coli growth and cell death assay*

Each sample of either MC4100(DE) or BL21(DE3) was added into a clear flat bottom 96-well plate (Corning CLS3997) and $OD_{600}$ was measured with a microplate reader (Infinite M200 Pro, Tecan). All samples were measured in triplicate. pH of each time points was measured with Hydrion pH paper (Micro Essential Laboratory, ranged from 4.5-7.5, 6.0-8.0, and 6.8-8.4). In order to test cell viability, samples from -3h, 0h, 2h, and 4h timepoints of time-course assay were resuspended in 170 mM NaCl and diluted to 0.01 $OD_{600}$ with DAPI (1 μg/ml). The cells were then analyzed by flow cytometry using a Sony SH800 cell sorter using a 488 nm laser and 450/50 FL1 (DAPI) and 665/30 FL4 (mPlum, for detecting TagRFP675) filters. The gating was created based on the fluorescence intensity of the control cells (Non-induced + Empty vector). To analyze and plot samples, 10,000 cells of each sample were counted and quantified using Cell Sorter Software (Version 1.7, LE-SH800 Series, Sony, San Jose, CA).

#### *Fluorescence vs $OD_{600}$ assay*

Samples from the fluorescence assay using LB plates were further used for comparing fluorescence with the $OD_{600}$ of cells. Briefly, samples were scraped from LB plates, and diluted into 0.1 $OD_{600}$ with 175 mM NaCl. The absorbance spectra in the range from 230 nm to 1,000 nm were measured with a microplate reader (Infinite M200 Pro, Tecan), using a clear bottom 96-well plate (Corning CLS3997). Purified TagRFP675 was added to the mock cells, MC4100(DE3) and BL21(DE3). Then the $OD_{600}$ of mock cells and the cells with purified TagRFP675 were compared with the induced cells which have a TagRFP675 expression vector, using a microplate reader (Infinite M200 Pro, Tecan). All experiments were performed in triplicate.
